# Molecular structure of the ESCRT III-based archaeal CdvAB cell division machinery

**DOI:** 10.1101/2025.09.15.676241

**Authors:** Tina Drobnič, Ralf Salzer, Tim Nierhaus, Margaret Jiang, Dom Bellini, Astrid Steindorf, Sonja-Verena Albers, Buzz Baum, Jan Löwe

## Abstract

Most prokaryotes divide using filaments of the tubulin-like FtsZ protein, while some archaea employ instead ESCRT-III-like proteins and their filaments for cell division and cytokinesis. The alternative archaeal system comprises Cdv proteins and is thought to bear some resemblance to ESCRT-III-based membrane remodelling in other domains of life, including eukaryotes, especially during abscission. Here we present biochemical, crystallographic and cryo-EM studies of the *Sulfolobus* Cdv machinery. CdvA, an early non-ESCRT component, adopts a PRC-domain/coiled-coil fold and polymerises into long double-stranded helical filaments, mainly via hydrophobic interfaces. Monomeric CdvB adopts the canonical ESCRT-III fold in both a closed and a distinct “semi-open” conformation. Soluble CdvB2 filaments are composed of subunits in the closed state, appearing to transition to the open, active state only when polymerised on membranes. Short N-terminal amphipathic helices in all CdvB paralogues, B, B1 and B2, mediate membrane binding and are required for liposome recruitment *in vitro*. We provide a molecular overview of archaeal ESCRT-III-based cytokinesis machinery, the definitive demonstration that CdvB proteins are *bona fide* ESCRT-III homologues and we reveal the molecular basis for membrane engagement. Thus, we illuminate conserved principles of ESCRT-mediated membrane remodelling and extend them to an anciently diverged archaeal lineage.

**Significance statement:** Membrane remodelling by ESCRT-III proteins is a fundamental and conserved process across the tree of life. The archaeal ESCRT-III-based cell division system (Cdv) drives cytokinesis in many archaeal groups, yet the molecular architecture of its components remained unknown, making it difficult to decipher the molecular mechanisms employed for cell division and cytokinesis in these organisms. We present structures of the Cdv machinery in *Sulfolobus* organisms that have been used previously to study the cell division process. We show that CdvA forms unexpected antiparallel helical filaments, while the ESCRT-III homologues retain the canonical fold we show they share with eukaryotic proteins. We demonstrate that N-terminal helices mediate membrane binding and that membrane contact, rather than polymerisation alone, likely triggers activation of Cdv ESCRT-IIIs.

## Introduction

Cell division culminates in cytokinesis, the physical separation of one cell into two. This involves dramatic membrane remodelling in all cells. The molecular machinery underpinning this process varies across life: bacteria and most archaea utilise FtsZ (tubulin)-based division machinery (1, 2), while eukaryotes use a contractile actomyosin ring and ESCRT-III (endosomal sorting complex required for transport III) proteins for cytokinesis (3, 4). In Crenarchaea, including S*ulfolobus*, cytokinesis is mediated by the ESCRT-III-like Cdv (cell division) system (5, 6).

In *Sulfolobus*, the Cdv system is composed of several division ring-forming components, CdvA, CdvB, CdvB1, and CdvB2, and a dedicated AAA+ ATPase CdvC (also called Vps4). It was first described in *Sulfolobus acidocaldarius*, with additional members identified in the related *Sulfolobus islandicus* (7). Division in *Sulfolobus* follows a defined sequence (Figure 1A). A ring of CdvA and CdvB first forms at the midcell. The ring functions as a non-contractile scaffold to recruit CdvB1 and CdvB2, forming a composite division ring. CdvB is then actively cleared from the ring by Vps4 and degraded by the proteasome. This allows CdvB1 and CdvB2 to remodel the membrane, driving constriction and abscission. The two proteins show a spatially patterned distribution in the dividing neck – CdvB1 is localised more peripherally, while CdvB2 is at the constricting membrane neck. There, it is believed to perform the final membrane abscission step, as evidenced by its strong loss-of-function phenotype (5, 8–10).

**Figure 1:**
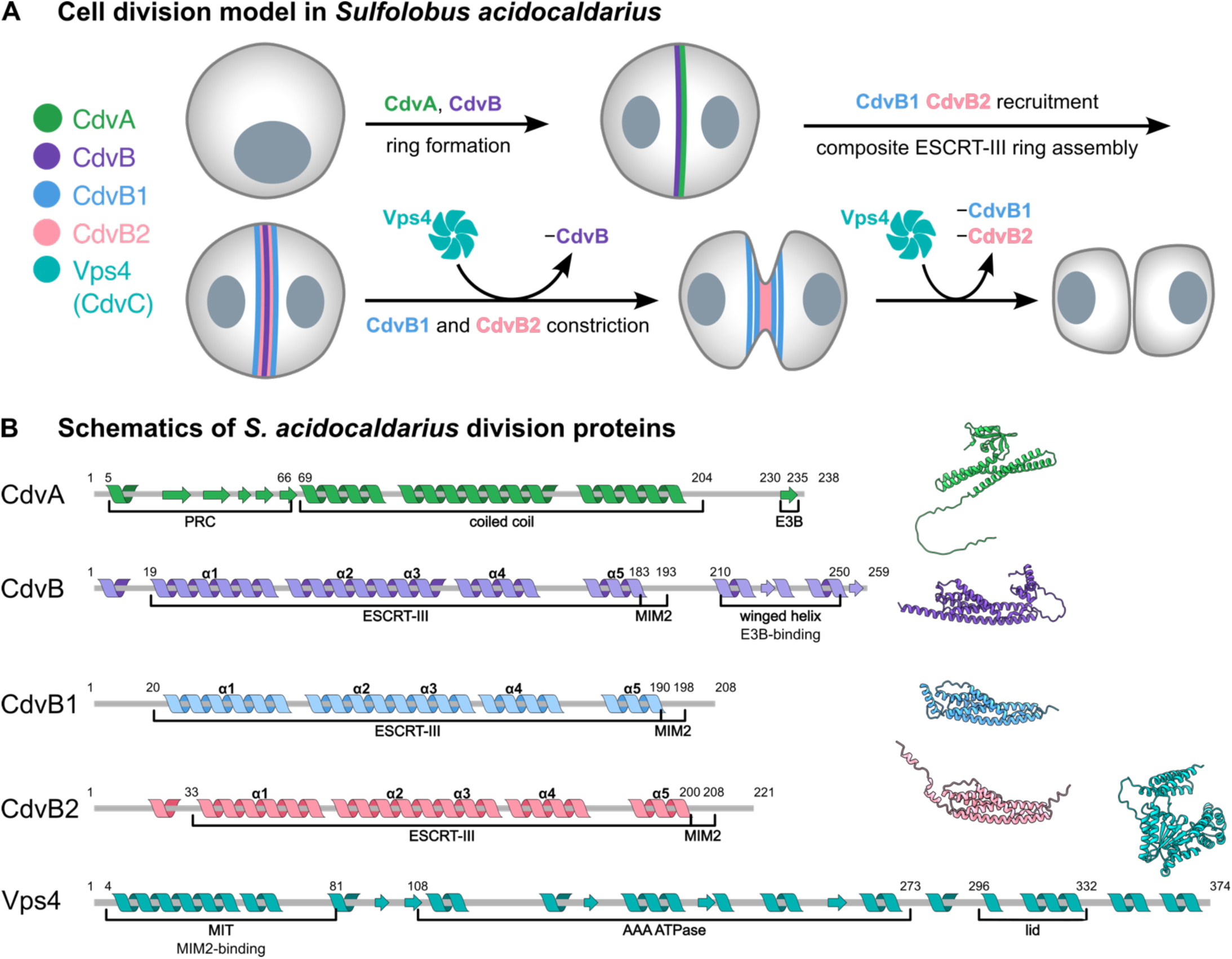
Archaeal Cdv cell division system. **(A)** Stepwise model of cell division in *Sulfolobus acidocaldarius*. Adapted from (1, 2). **(B)** Schematics of secondary structure and domain organisation of *Sulfolobus* Cdv cell division proteins. Models of *S. acidocaldarius* proteins from the AlphaFold2 database (3) are shown in corresponding colours on the right. Helices α1-α5 are denoted in ESCRT-III proteins. Residue numbers refer to *S. acidocaldarius* sequences, secondary structure was predicted with Jpred 4 (4).

CdvA is an archaeal protein unique to the Cdv system (11). It is predicted to have a PRC domain, a coiled-coil domain, and a C-terminal E3B (ESCRT-III-binding) sequence. Though it lacks homology to any ESCRT component, it is essential for templating downstream ESCRT-III assembly (12) via the E3B peptide directly binding CdvB (8).

Though lacking the homologous upstream components, the Cdv system is related to eukaryotic ESCRT-III membrane remodelling machineries as determined by amino acid sequence comparisons. Canonically, the ESCRT-III functional core is made of five α-helices (α1-α5). Membrane remodelling is driven by polymerisation of ESCRT-III proteins that transition from a “closed” monomeric (13, 14) to an “open” polymeric conformation (15–17). CdvB, CdvB1, and CdvB2 are believed to be ESCRT-III homologues. All are predicted to share the characteristic five-helix ESCRT-III domain and a MIM2 motif for interaction with Vps4. (Figure 1B). The three proteins are paralogues, likely having arisen via two gene duplication events from an ancestral CdvB (18).

Despite the central role of the Cdv system in division, high-resolution structures exist in the Protein Data Bank (PDB) only for the ATPase CdvC (19, 20). Thus, the molecular mechanisms underlying Cdv function remain largely unclear. Here, we report the molecular structures of components of the *Sulfolobus* Cdv system: CdvA, CdvB, and CdvB2. We show that CdvA adopts a novel fold and polymerises into long antiparallel helical filaments via hydrophobic interactions. The structures of monomeric CdvB and polymerised CdvB2 reveal canonical ESCRT-III features. Unexpectedly, a CdvB2 filament in solution is composed of subunits in the closed state and appears to transition to the open conformation only when polymerised on a membrane. As with CdvB and CdvB1, this membrane interaction is mediated by short N-terminal amphipathic α-helices.

## Results

### Constitutive helical filaments of CdvA are stabilised via hydrophobic interactions

CdvA is unique to archaeal ESCRT-III systems, with no clear eukaryotic homologues. In *Sulfolobus*, it is the first Cdv component recruited to midcell, where it forms a ring and templates assembly of subsequent ESCRT-III rings (12).

CdvA from *Metallosphaera sedula* has been shown to form double stranded filaments that were reportedly stabilised by DNA (11). To investigate how the protein polymerises, we purified full-length *S. islandicus* CdvA for structural analysis. It formed filaments on its own, which only dissolved in 1M CHES buffer at pH 11. As CdvA is predicted to have a flexible C-terminal linker, we removed the linker and the E3B peptide region to aid in crystallisation (CdvA^ΔC^, residues 1-205). Removing the trailing 33 residues did not disrupt polymerisation (Figure 2A) and enabled us to determine the molecular structure of crystallised CdvA^ΔC^ at 2.9 Å resolution. Interestingly, CdvA^ΔC^ formed a two-stranded helical filament with anti-parallel strands in the crystal (Figure 2B). A dimer of CdvA is the repeating unit in the filament (dark green and magenta chains). The N-terminal PRC domain of each CdvA^ΔC^ molecule sits on the inside of the helix, while the three-helix coiled-coil domain faces the outside.

**Figure 2:**
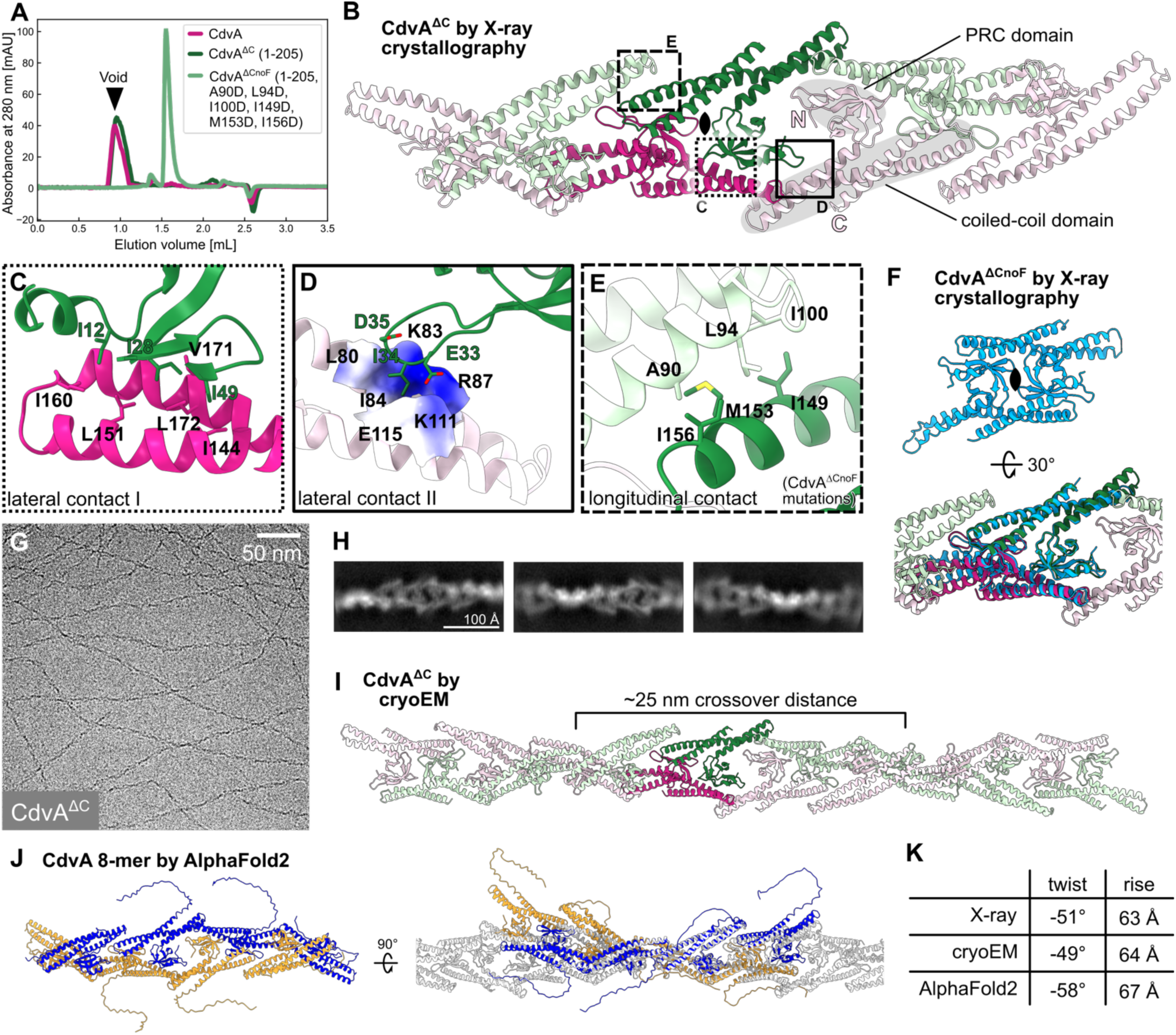
CdvA forms constitutive double-stranded helical filaments. **(A)** Size exclusion chromatography profiles of purified CdvA constructs. Filament formation is abolished by mutating six hydrophobic residues (CdvA^ΔCnoF^). **(B)** *S. islandicus* CdvA^ΔC^ as determined by X-ray crystallography forms a dimer (darker colour), which repeats to form a double-stranded helical filament in the protein crystal. One strand (protofilament) is coloured green, the other pink. Each CdvA subunit has a PRC domain and a three-helix coiled-coil domain. **(C)** Hydrophobic and **(D)** hydrophilic lateral contacts between CdvA^ΔC^ protofilaments. View is rotated by 180° relative to panel B. **(E)** Longitudinal contacts within each protofilament are hydrophobic interactions between coiled-coil domains. Mutating these six residues prevents filament formation (panel A). **(F)** A The non-polymerising CdvA mutant (CdvA^ΔCnoF^) forms a dimer similar to CdvA^ΔC^ as determined by X-ray crystallography. Bottom panel: overlaid with CdvA^ΔC^ structure from panel B. **(G)** Representative cryo-electron micrograph of CdvA^ΔC^ filaments. **(H)** 2D class averages of CdvA^ΔC^ filaments. **(I)** Cryo-EM structure of CdvA^ΔC^ agrees with the double-stranded helical filament determined by X-ray crystallography. **(J)** AlphaFold2 prediction of eight CdvA chains agrees with experimentally determined CdvA^ΔC^ filament structures. Overlaid on the cryo-EM structure (grey) in the right panel. **(K)** Table of approximate helical parameters of the crystal, cryo-EM, and AlphaFold2 structures. For crystal and AlphaFold2 models, parameters were calculated with *relion_helix_toolbox* on simulated maps (*molmap* command on PRC and coiled-coil domains in ChimeraX).

Inspecting the filament structure revealed that lateral interactions between the two strands involve a PRC domain on one strand contacting the coiled-coil domain of two subunits on the opposite strand. Lateral contact I is mediated by extensive hydrophobic interactions (Figure 2C), while lateral contact II consists of three hydrophobic residues nested in an otherwise electrostatic pocket (Figure 2D). Within each protofilament (strand), the coiled-coil domains contact one another in a head-to-tail manner. This interface is made up of six hydrophobic residues, three on each subunit (Figure 2E). To test the role of this longitudinal interaction on CdvA’s ability to polymerise, we mutated the six positions to aspartate residues (A90D, L94D, I100D, I149D, M153D, I156D). This resulted in a non-polymerising CdvA^ΔCnoF^ protein (Figure 2A). A 2.2 Å structure of CdvA^ΔCnoF^ confirmed that the mutations did not substantially alter the CdvA fold. The protein crystallised as a dimer that closely matched the dimeric repeat unit of the CdvA^ΔC^ filament (Figure 2F), but did not itself form a filament in the crystal. We aligned the dimers using the PyMOL (v2.5) *align* command. The reported backbone RMSD was 1.56 Å after outlier rejection, reflecting conservation of the overall fold. Taken together, the hydrophobic coiled-coil-to-coiled-coil interface appears to be essential for CdvA filament formation.

To verify the CdvA filament observed in the protein crystals we turned to cryo-EM (Figure 2G, H). The final helical reconstruction at 4.1 Å resolution (Figure 2I) was very similar to the filament observed in the crystals, having the same subunit arrangement and almost identical helical parameters. A similar polymer assembly was also recapitulated by AlphaFold2 predictions (Figure 2J), albeit with somewhat different calculated helical parameters (Figure 2K). However, the model retained the same fold, subunit organisation, longitudinal and lateral interfaces as the experimental structures. Taken together, these data confirm that CdvA polymerises into double-stranded, antiparallel, helical filaments with strong hydrophobic interaction surfaces.

### Archaeal CdvB has an ESCRT-III fold

In analysing the ESCRT-III homologues involved in cytokinesis in Sulfolobales, we focussed initially on CdvB, the first ESCRT-III to be recruited to the site of division. The *S. islandicus* homologue was truncated to remove the C-terminal linker and winged helix domain (residues 193-259), to aid crystallisation, and two additional methionine residues were introduced to enable crystallography phasing with selenomethionine (I69M, I125M). A structure of CdvB was determined at 2.6 Å (P 4_1_ 2_1_ 2), which revealed a canonical ESCRT-III fold of 5 α-helices in the “closed” conformation as reported for other ESCRT-III proteins (Figure 3A). In most previously reported cases (with the notable exception of IST1 (17)), ESCRT-III proteins exist in a closed conformation when monomeric, but transition to the open form once polymerised (13–16). A second CdvB crystal form (P 2_1_ 2_1_ 2_1_) revealed an alternate conformation at 2.2 Å resolution (Figure 3B). Here, helices α1-α4 remain largely unchanged, but helix α5 is swung open approximately 120°, making it no longer folded over the α1-α2 hairpin. We term this conformation “semi-open” as only helix α5 changes its relative position while the rest remain as in the closed conformation.

**Figure 3:**
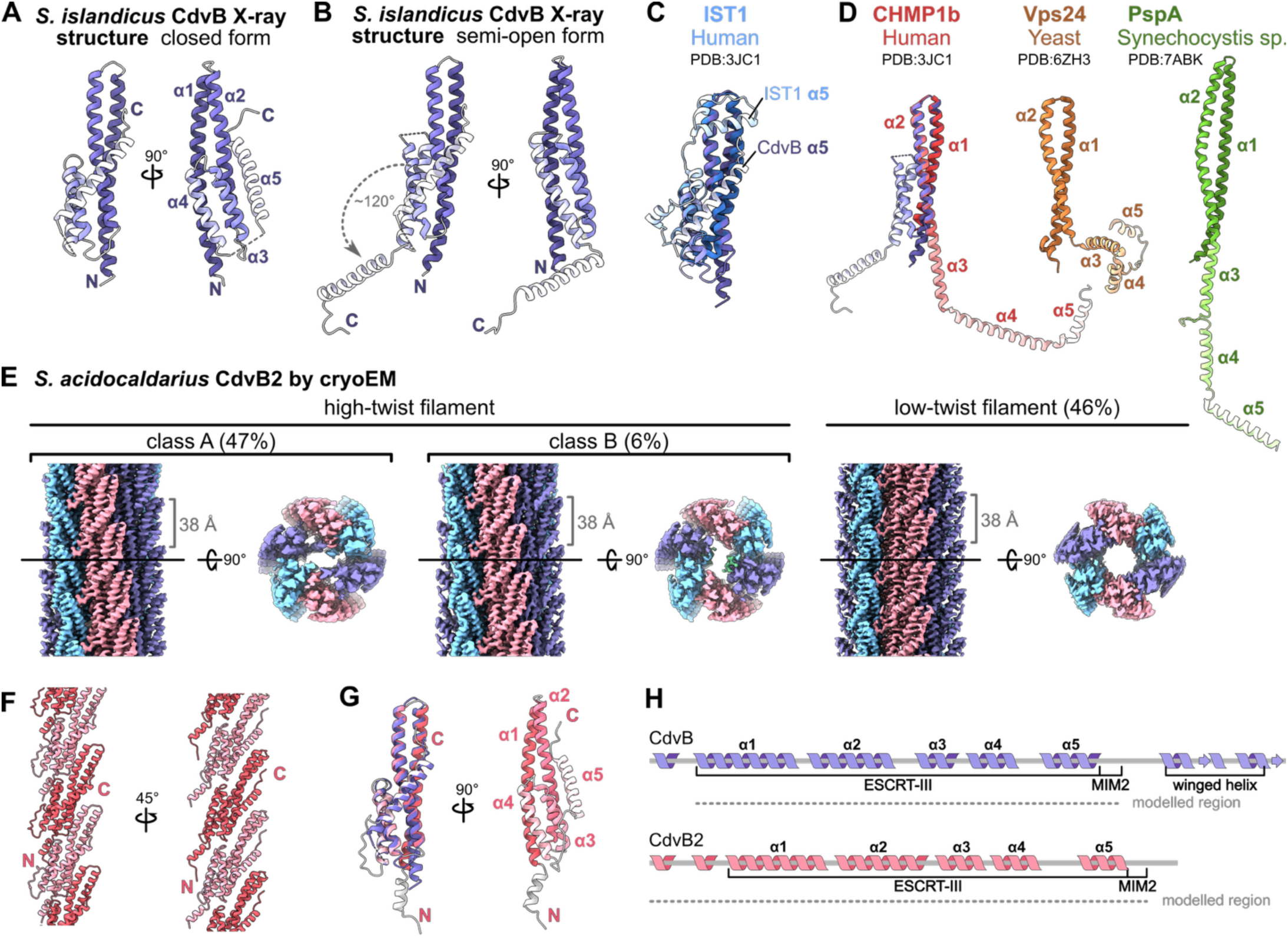
Structures of archaeal CdvB and CdvB2 ESCRT-IIIs. **(A)** *S. islandicus* CdvB ESCRT-III protein in the closed conformation as determined by X-ray crystallography (P 41 21 2). Helices 1-5 are coloured in increasingly lighter shades. **(B)** A second CdvB crystal form (P 21 21 21) shows a semi-open conformation, where helix 5 swings open by ∼120°. Closed conformation of helix 5 shown transparent. **(C)** Comparison of CdvB and IST closed-form structures. **(D)** Comparison of the semi-open form of CdvB with different open-form ESCRT-IIIs: CHMP1b, Vps24, and PspA. **(E)** *S. acidocaldarius* CdvB2 forms helical filaments of six protofilaments. Three filament conformations fall into two main classes, high-twist and low-twist. Side view and central slices along the XY and YZ axes are shown for each. Inter-subunit spacing within a protofilament (∼38 Å) is consistent across the three classes. **(F)** Atomic model of a low-twist protofilament. N-termini face the filament lumen, C-termini are surface-exposed. **(G)** Atomic model of a low-twist CdvB2 monomer, in closed form. Left panel: aligned to *S. islandicus* CdvB from panel A (purple). 1.13 Å RMSD after outlier rejection, aligned with ChimeraX *matchmaker*. **(H)** Cartoon schematics of CdvB and CdvB2, updated to reflect experimentally determined secondary structures of the modelled regions.

We compared these CdvB monomer structures with ESCRT-III structures from organisms in other domains of life determined previously. Superposition of the closed-form CdvB and human IST1 highlights the ESCRT-III fold being conserved between archaea and eukaryotes (Figure 3C). The main difference between the two is the position of helix α5, which sits lower on the α1-2 hairpin in CdvB. Comparing the semi-open CdvB structure with a range of different ESCRT-III structures reveals that CdvB can adopt a distinct conformation (Figure 3D). In open-form ESCRT-III proteins, from bacteria to humans, helices α3-5 unfurl, with α3 becoming a continuous extension of α2. In the semi-open conformation of CdvB, helices α3-4 remain packed to the side of the α1-2 hairpin. A different “extended” ESCRT-III conformation was previously described in yeast Vps24 filaments (21) (Figure 3D, ochre), where helices α3-5 move away from the α1-2 hairpin, but are not fully extended.

### A closed-conformation CdvB2 filament buries its amphipathic helices

ESCRT-III monomers can form polymers that bind to and remodel membranes. To further investigate archaeal Cdv polymerisation, we sought to structurally characterise CdvB polymers. Polymerisation could be induced by the presence of either CHAPS or CHAPSO, or lipid monolayers. However, the samples were not amenable to high-resolution cryo-EM due to bundling and a lack of helical twist, leading to preferred orientation (Figure S2ABC).

We instead turned to CdvB2, which is thought to perform the final membrane abscission during cytokinesis, in the better-characterised *S. acidocaldarius* system (10). Adjusting the buffer pH to 6 triggered the formation of CdvB2 filaments, which were then imaged by cryo-EM (Figure S3A).

We solved the structure of *S. acidocaldarius* CdvB2 filaments in three different forms (Figure 3E). 2D class averages revealed two main filament types, high-twist and low-twist (Figure S3B). The high-twist filaments were further separated into two classes (A and B). All three filament types consisted of six protofilaments with longitudinal subunit spacings of ∼38 Å and differed slightly in their helical parameters. The primary difference between high-twist classes A and B was in their lumenal densities. Atomic modelling revealed that, in all filament forms, the N-termini face the lumen of the filaments, while the C-termini face outwards. This is consistent with the C-terminal MIM2 motif being solvent-exposed and accessible for remodelling by Vps4. In all cases, we saw density for only the first few amino acids of the MIM2 motif, indicating flexibility.

As in the case of CdvB, CdvB2 also adopts the characteristic 5-helix ESCRT-III fold. Moreover, the structures of individual CdvB2 and CdvB chains are strikingly similar, despite different source organisms and oligomeric states (*matchmaker* in ChimeraX (v1.7) reported RMSD of 1.13 Å after outlier rejection, Figure 3G, left panel). However, in all three filament forms, the CdvB2 subunits were found in a closed conformation (Figure 3FG), rather than in an open conformation as would be expected for a polymerised ESCRT-III protein. While truncated IST1 has been reported to generate polymers in a closed form as part of a composite polymer, to our knowledge this is the first example of an ESCRT-III assembling into a closed-form homopolymer.

One of the main differences that distinguishes the monomer structure from the 3 types of CdvB2 filament is the N-terminal region, which adopts different conformations (Figure 4A). In all filament cases, hydrophobic residues consistently pack together and are shielded from solvent in the lumen. The hydrophobic residues tucked away on the inside of the filaments clustered in three regions, which were short α-helices with a hydrophobic face. The first two regions (M1-W11 and I19-F27) are short amphipathic α-helices (named here amph1 and amph2), while the third (P32-Y42, amph3) overlaps with the start of helix α1 of the ESCRT-III domain, which forms the coiled-coil hairpin with α2 (Figure 4B). This third helix therefore has two hydrophobic faces: the coiled-coil interface facing α2, and the outer hydrophobic face that is involved in N-terminal hydrophobic interactions. Thus, CdvB2 has N-terminal amphipathic helices which pack in different ways to occlude their hydrophobic regions from solvent. This analysis led us to inspect the N-terminal regions of CdvB, CdvB1, and CdvB2 of different *Sulfolobales* species to assess the conservation of their N-terminal amphipathic regions (Figure 4C).

**Figure 4:**
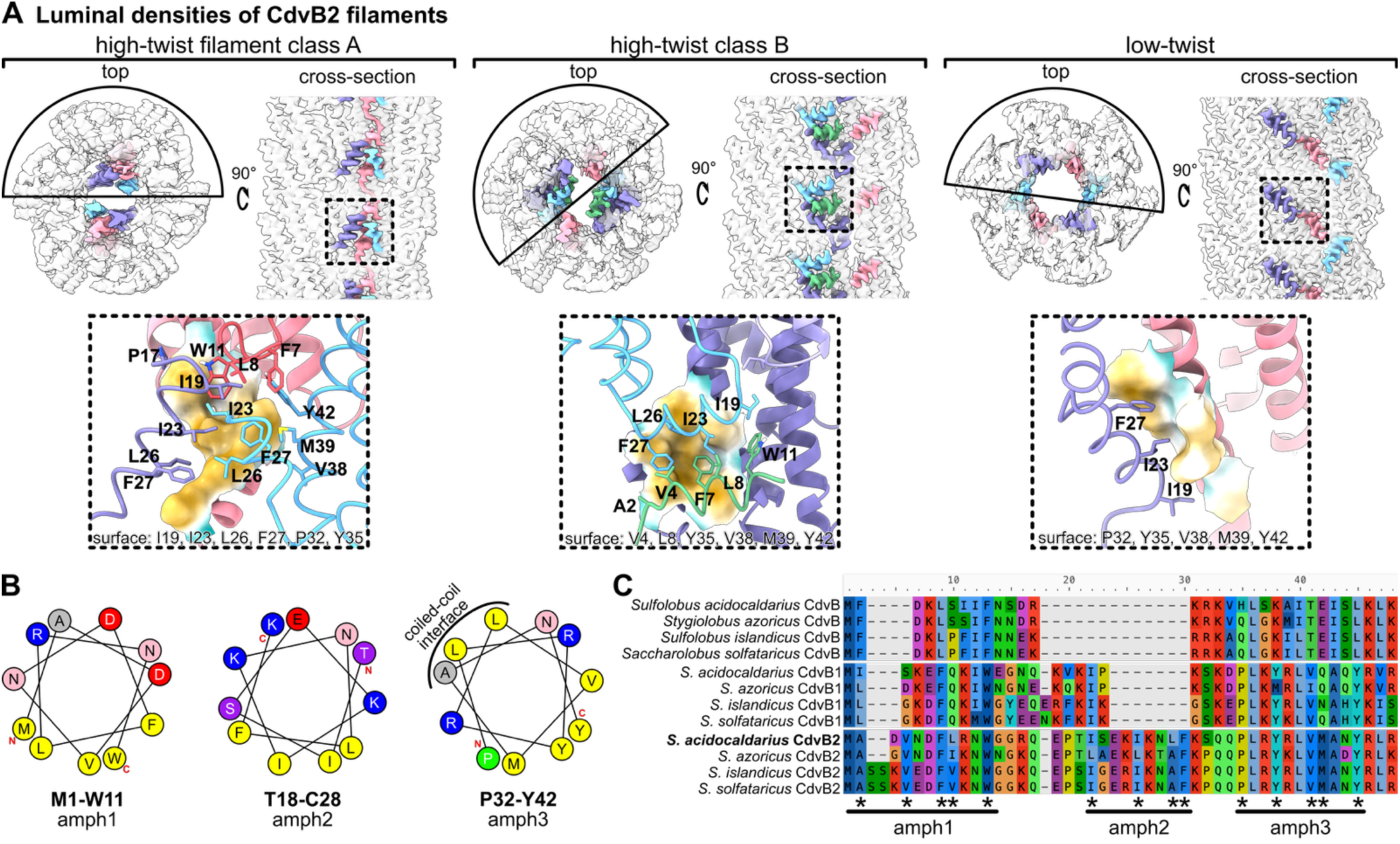
CdvB2 N-terminal region has amphipathic helices. **(A)** Top row: cryo-EM maps of CdvB2 highlighting the different arrangements of N-termini (in colour) on otherwise transparent maps. Bottom row: cryo-EM derived atomic models of CdvB2 N-termini, corresponding to regions in dashed boxes in the top row. **(B)** Helical wheel diagrams of three α-helical regions involved in hydrophobic interactions in panel A, demonstrating they are short amphipathic helices. Calculated with the HeliQuest server (5). **(C)** Sequence alignment of N-terminal regions of CdvB, CdvB1, and CdvB2 from related *Sulfolobales*. Hydrophobic residues from panels A, B are highlighted with asterisks. Aligned with MAFFT online in auto mode (6).

First, it was important to ensure that the predicted start codon is correct in each case. Initial sequence alignments of CdvB1 sequences indicated that the start codons of some current database entries may be mis-assigned (relevant here: WP_014512765.1 for *S. islandicus*, WP_011277365.1 for *S. acidocaldarius*) - as they have been predicted to have long N-terminal regions that are not present in other CdvB1s, and contain a methionine at the position that corresponds to the first residue of other CdvB1s (Figure S2D). The long leading N-terminal sequence of *S. islandicus* CdvB1 has three possible start codon positions (M1, M11, and M46). To assess which start codon is used *in vivo*, we expressed the three CdvB1 variants (aa 1-153, 11-153, 46-253) in *E. coli* and visualised the proteins in lysates by Western blotting using an anti-SaciCdvB1 antibody (10). We compared *E. coli* lysates with the overexpressed proteins with *S. islandicus* and *S. acidocaldarius* cell lysates (Figure S2E). CdvB1^46-253^ corresponded well with the apparent size of the native *S. islandicus* CdvB1 band, indicating that the start codon is indeed mis-annotated in at least some database entries. Throughout this manuscript, we therefore trimmed the leading N-terminal sequence and renumber residues based on the re-assigned starting methionine.

Having determined the N-terminus, we could then compare regions. The N-terminal regions of these proteins have a high level of sequence diversity compared to the core ESCRT-III domain. Nevertheless, the residues forming the hydrophobic faces of the three amphipathic helices are conserved in CdvB2 orthologs. CdvB and CdvB1 sequences maintain some of the hydrophobic residues in amph1 and amph3 but lack the corresponding amph2 sequence. Helical wheel diagrams of amph1 and amph3 in *S. acidocaldarius* CdvB and CdvB1 indicate that these regions could form short amphipathic helices - similar to those observed in CdvB2 (Figure S3G).

### Interactions between CdvB paralogues and membranes

CdvB, CdvB1, and CdvB2 are believed to polymerise on archaeal membranes and remodel them during cell division, driving cytokinesis. To understand this process better, we assayed how the three CdvB paralogues associate with different membranes *in vitro*.

We incubated *S. islandicus* CdvB, CdvB1 (with the reassigned start codon) and CdvB2 in different combinations together with lipid nanotubes (LNTs) composed of *E. coli* total lipid extract and galactosyl(β)ceramide. Looking at the samples with cryo-EM (Figure 5A), both CdvB and CdvB2 coated the surface of the membrane tube in a relatively regular manner. Under the same conditions, we could not see any CdvB1 bound to LNTs. Mixing all three paralogues and incubating them with LNTs produces a single-layer protein coat, although we could not determine whether it is made of a single CdvB paralogue or a mixture of all three. When the dedicated ATPase Vps4 was added to the mix and supplemented with ATP, the assembled protein coat was much thicker and less regular.

**Figure 5:**
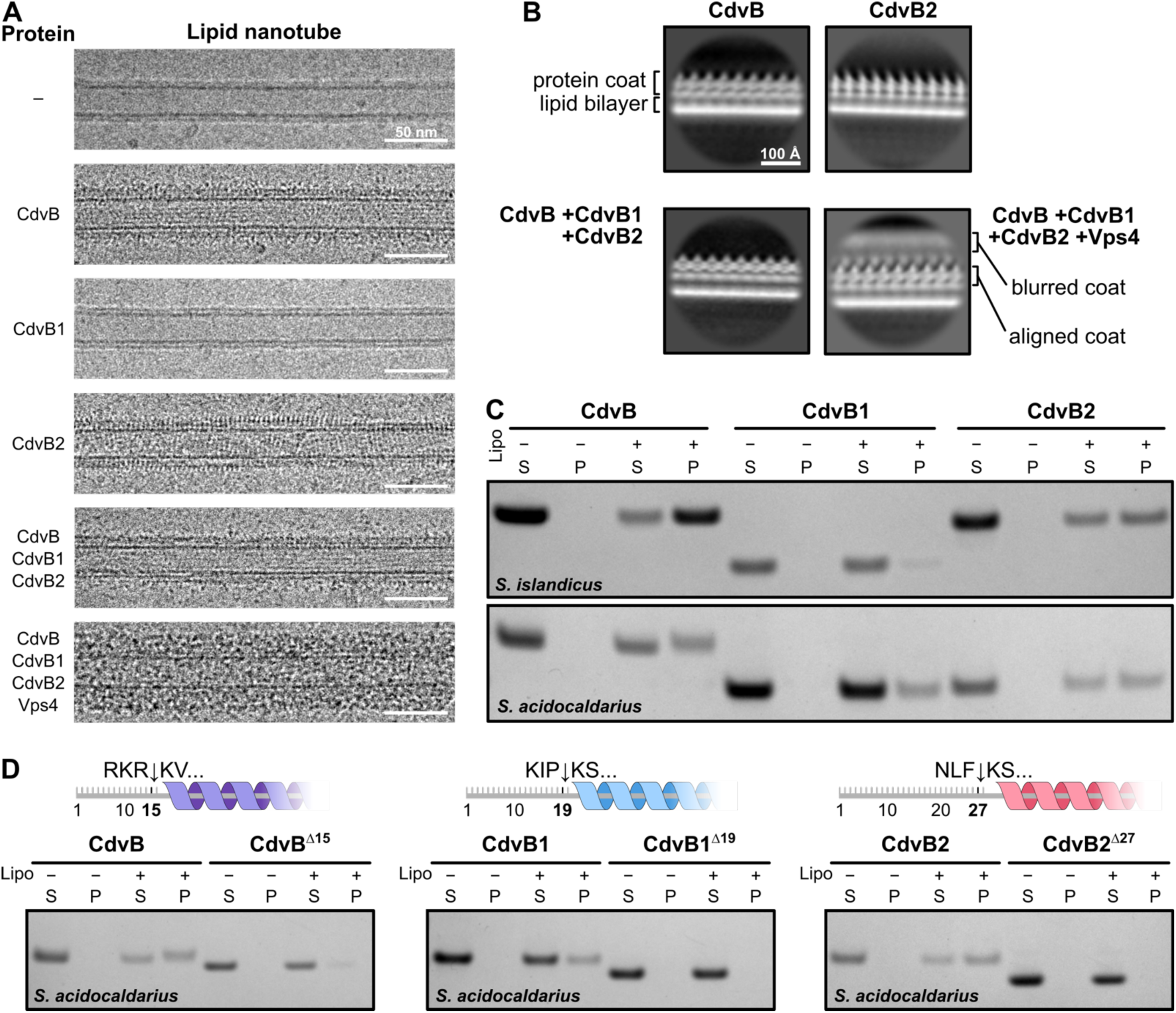
Interactions between CdvB-family proteins and lipid membranes. **(A)** Cryo-EM micrographs showing lipid nanotubes (LNTs) in the presence of different combinations of *S. islandicus* Cdv proteins. **(B)** 2D class averages of LNTs mixed with different *S. islandicus* Cdv proteins from panel A. **(C)** SDS-PAGE gels of *S. acidocaldarius* and *S. islandicus* CdvB, CdvB1, and CdvB2 pelleting in the presence and absence of archaeal liposomes. (S: supernatant. P: pellet). **(D)** SDS-PAGE gels comparing pelleting with archaeal liposomes of full-length and N-terminally truncated *S. acidocaldarius* CdvB, CdvB1, and CdvB2. Positions of truncations are indicated with arrows in cartoon schematics.

To better resolve the structures of these protein coats, we performed 2D classification and averaging of the coated nanotube edges (Figure 5B). CdvB, CdvB2, and the CdvB+CdvB1+CdvB2 condition all form single-layered protein coats with ∼38 Å repeats along the LNT’s long axis. The same spacing has been reported above in the monolayer-bound *S. islandicus* CdvB and soluble *S. acidocaldarius* CdvB2 filaments (Figure S2AC, Figure 3E). The multi-layered coat resulting from addition of Vps4 is irregular as mentioned, the 2D average showing a blurred protein coat on top of the aligned 38 Å-repeat protein layer. It remains unclear whether the additional density represents Vps4 binding to exposed MIM2 motifs, or whether it is a multi-layered ESCRT-III coat that requires Vps4-mediated remodelling for assembly.

Unlike most lipids of bacteria and eukaryotes, which contain fatty acids ester-linked to a glycerol backbone, archaeal lipids contain ether-linked isoprenoid chains. Membranes of thermophiles such as *Sulfolobus* additionally have a high proportion of membrane spanning tetraether lipids, which likely confer greater thermal stability on the membrane (22). Since LNTs were made with bacterial lipids, we wanted to evaluate whether archaeal lipids alter CdvB paralogue membrane recruitment.

We generated archaeal liposomes using lipids extracted from *S. acidocaldarius* and used them to perform liposome pelleting assays. After an initial spin that removed protein aggregates and any pre-formed filaments, we incubated the soluble proteins with archaeal liposomes for 15 min and pelleted the liposomes, including any associated protein. We performed the experiment using both *S. islandicus* and S*. acidocaldarius* proteins.

The results of these pelleting assays were effectively consistent with LNT binding. CdvB and CdvB2 efficiently pelleted with archaeal liposomes, while CdvB1 remained mainly in the soluble fraction (Figure 5C). This was consistent across the two species. The fact that these proteins exhibit the same binding profile in presence of *E. coli* and *S. acidocaldarius* lipids indicates that the CdvB paralogues are not selective for lipid backbones or tails when polymerising on membrane.

### N-terminal amphipathic helices are required for efficient membrane recruitment

N-terminal amphipathic helices have shown to be required for membrane binding and remodelling in eukaryotic ESCRT-IIIs (23, 24). To probe how CdvB proteins interact with the membrane, we tested the role of the N-terminal amphipathic helices identified above (Figure 4, Figure S3G). Having established an assay for CdvB membrane recruitment, we assessed the role of the N-terminal amphipathic helices in this process.

As we initially found short amphipathic helices in the *S. acidocaldarius* CdvB2 structure, we performed this experiment with *S. acidocaldarius* proteins as a model. We truncated the first 15 residues of CdvB (CdvB^Δ15^), 19 in CdvB1 (CdvB1^Δ19^) and 27 residues of CdvB2 (CdvB2^Δ27^), removing amph1 and amph2 (amph3 of CdvB2 was left intact because it corresponds with the start of the core ESCRT-III domain). The truncated proteins were then tested in the membrane binding assay. In each case, the N-terminal truncation exhibited a drastic reduction in liposome binding (Figure 5D). While CdvB1^Δ19^still showed a faint band in the pellet fraction, membrane binding was completely abolished in CdvB^Δ15^ and CdvB2^Δ27^.

This indicates that the N-terminal regions are essential for CdvB paralogues to associate with membranes.

### CdvB2 on lipid nanotubes

To assess whether full length CdvB2 polymers undergo a conformational change upon membrane binding, we prepared LNTs incubated with *S. acidocaldarius* CdvB2 and imaged them by cryo-EM (Figure S4A). To inspect the fine structure of the coat, we performed 2D classification on protein-coated LNT edges. The classes showed spiked edges (Figure 6A, Figure S4B), similar to the *S. islandicus* samples. To assess whether *S. acidocaldarius* CdvB2 adopts a closed or open conformation in this context, we generated simulated 2D projections of closed and open CdvB2 filaments based on our cryo-EM structure and AlphaFold2 (25, 26), respectively. When directly compared, experimental 2D classes of the protein edge more closely match the simulations in the open conformation (Figure 6A).

**Figure 6:**
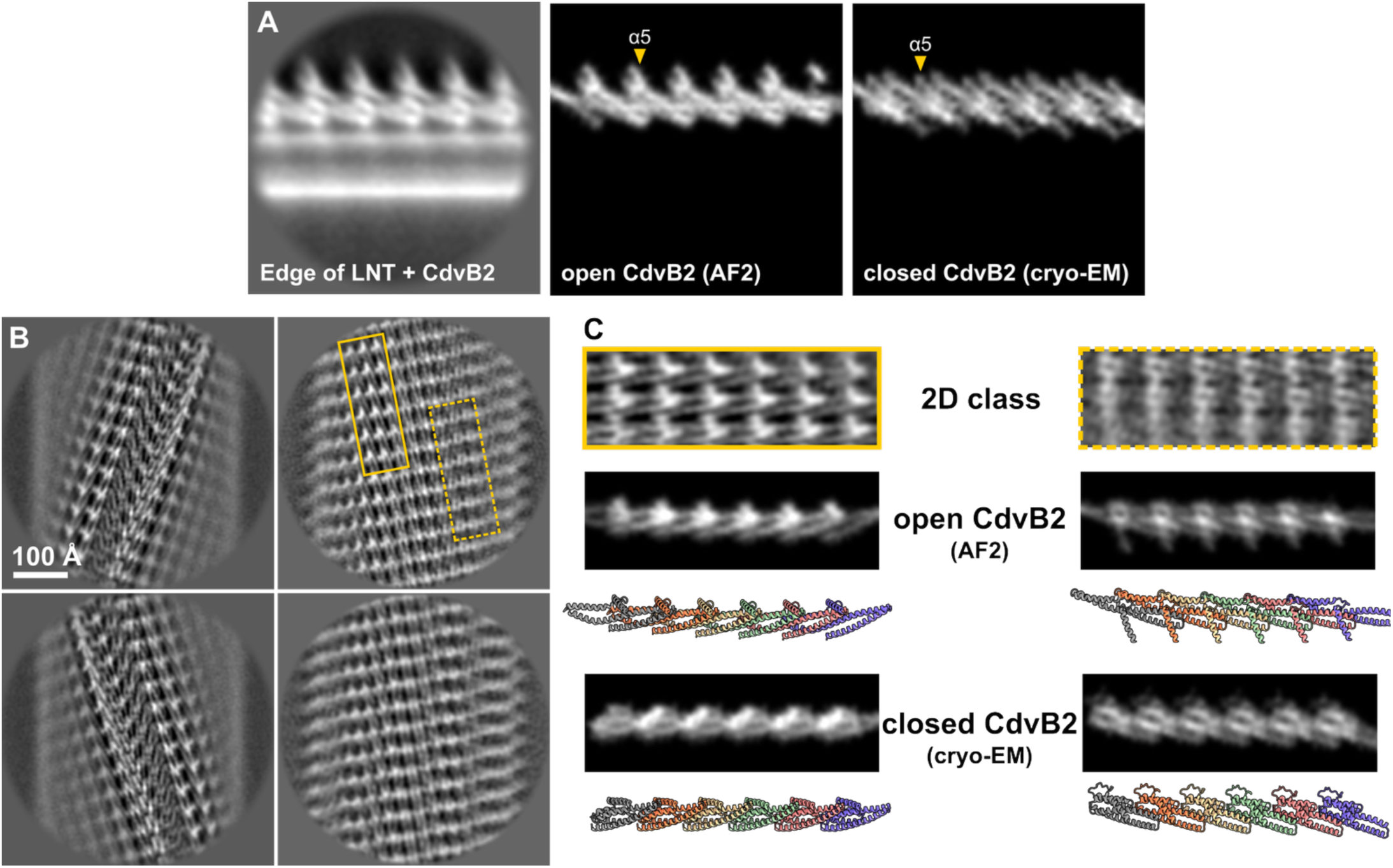
Membrane-bound CdvB2. **(A)** 2D class average of LNTs coated with *S. acidocaldarius* CdvB2 (left) alongside simulated 2D projections of open (modelled by AF2) and closed (cryo-EM, this work) CdvB2 filaments. To simulate projections, a density map at 10 Å resolution was generated from an atomic model with ChimeraX *molmap* and projected using relion_project with added white noise. **(B)** 2D class averages of LNTs coated with CdvB2. Membrane signal has been subtracted. The two main types of high-resolution 2D classes contained seams (left column) or continuous CdvB2 coats (right column). **(C)** Two regions of the continuous protein coat from (B) are compared with simulated 2D projections of open (middle) and closed (bottom) CdvB2 filaments. Corresponding molecular models are shown for each simulated projection.

To better assess the CdvB2 conformation in the context of a membrane bound polymer, we processed a second dataset of CdvB2 on LNTs, treating the whole tube as a filament. To improve alignment we subtracted the dominating membrane signal from micrographs, as implemented in (27), borrowed from the field of microtubule-associated proteins (28). This greatly improved protein alignment (Figure S4G), enabling us to resolve 2D classes to secondary structure resolution (Figure 6B). Two main types emerged: some had a continuous protein coat while others had a prominent seam. To further assess the conformation of the membrane-bound CdvB2, we compared different regions of the tubular protein array with simulated 2D projections of closed- and open-form filaments from equivalent angles (Figure 6C). The experimental averages appeared more similar to the open CdvB2 model than the closed-form filament. These data reinforced the conclusion that membrane binding triggers CdvB2 to polymerise in the open conformation.

We noted that in some of these images we observed non-membrane bound CdvB2 filaments in the background. All were low-twist filaments with monomers in the closed conformation, indistinguishable from the low-twist CdvB2 structure solved earlier (Figure S4C-F). This implies that polymerisation on membrane induces the protein to undergo a conformational switch from the closed to the open form.

## Discussion

In this work, we provide a structural and biochemical framework for understanding Cdv-based cell division. We reveal how elements of the core machinery assemble into polymers and how they engage with membranes to drive division.

CdvA, the non-ESCRT component, has a fold containing a PRC and three-helix coiled-coil domain. It forms antiparallel double-stranded helical filaments built from dimeric subunits and stabilised by hydrophobic interactions between the coiled-coil domains. This double filament architecture is consistent with earlier observations of *M. sedula* CdvA by negative stain electron microscopy (11, 29). While it was previously reported that *M. sedula* CdvA requires DNA for polymerisation, we did not observe DNA in our cryo-EM sample or in the final polymer structure. The positioning of PRC domains in lateral cross-strand interactions provides a structural explanation for previous observations that CdvA lacking this domain fails to polymerise (30). Although CdvA is thought to associate with the membrane in cells (8, 12), CdvA filaments do not show *in vitro* membrane binding capacity (29), and we did not observe putative membrane binding regions on the surface of the filament. Thus, we speculate that CdvA may require additional, yet-to-be-identified partners that activate it or tether it to membranes *in vivo*.

The helical nature of the CdvA filament seems at odds with its function as a ring forming protein in the cell, which could be assumed to require association with the membrane with one surface. Possible ways out of this conundrum are: the filament re-assembles to form an alternate structure when membrane-bound (as observed here for CdvB2); the membrane attachments are not continuous but periodic and occur where the helical twist allows it, reducing the number of attachment points; or CdvA filaments do not assemble into a complete ring but instead form short discontinuous filaments, similar to FtsA and MreB which act in bacterial cell division and elongation, respectively (31). Further studies will be needed to unravel the enigmatic role of this protein in cell division and cytokinetic ring initiation.

Through our analysis of the main ESCRT-III proteins in Sulfolobales, we found that CdvB and CdvB2 adopt the canonical ESCRT-III fold, a result that experimentally confirms their evolutionary and functional placement within the ESCRT-III protein superfamily (32).

Interestingly, however, our monomeric CdvB structures reveal both a closed and a previously undescribed semi-open conformation, suggesting a possible conformational intermediate along the transition to fully open, polymer-forming state seen in other ESCRT-IIIs. This makes clear the flexibility of ESCRT-III polymers, which likely enables them to play a unique role in the deformation and scission of membranes.

We show that soluble, lipid-free CdvB2 filaments are composed entirely of subunits in the closed state, a unique example of a full-length ESCRT-III filament adopting this conformation in the context of a homopolymer. A truncated eukaryotic ESCRT-III protein, IST1, was previously shown to assume a closed conformation when co-polymerised with CHMP1B, or when mutated (17, 33). This led to the suggestion that some ESCRT-IIIs may function solely in the closed state. In our experiments, however, the presence of membranes was sufficient to induce the conformational switch to the open state. Thus, while CdvB2 subunits assembled in a closed state in soluble filaments, they adopted an open state when polymerised on a lipid membrane. This challenges a model in which polymerisation alone triggers the closed-to-open switch. Instead, it appears that polymerisation of CdvB proteins on a membrane triggers a change in conformation.

Our analysis also revealed that the short N-terminal amphipathic helices in CdvB paralogues mediate membrane binding and are essential for recruitment *in vitro*. Amphipathic helices are common features of many membrane-associated proteins (34). They have also been observed in both in eukaryotic (16, 23, 24) and bacterial ESCRTs (35–38), where they bind membrane and are essential for proper function. In *Sulfolobus*, these amphipathic regions are notably short (9-11 residues) and, in the case of CdvB2, there are two of them. Despite their size, they are clearly sufficient to mediate membrane binding, likely aided by avidity effects in the polymer form.

In cells, CdvA and the three CdvB paralogues studied here have been reported to be recruited sequentially and to localise to distinct regions of the constricting division neck (10). While all three CdvB proteins have N-terminal membrane-targeting regions, these segments of sequence are the most diverse across paralogues. These sequence differences may contribute to their differential recruitment, distinct functions, membrane affinities, or localisation on the dividing cell membrane (10).

Future research should focus on the mechanism of membrane remodelling, using both *in vitro* and *in vivo* approaches. Reconstituting the constriction machinery on the inside of liposomes and imaging division rings with fluorescence microscopy and cryo-EM would reconcile dynamic and structural information about the mechanism of ESCRT-III dependent cell division. In addition, it will be important to image cell bridges by cryo-electron tomography (cryo-ET) (10) at different stages in the division process to assess the structure of these filaments in their native context. For this we need mutants that will trap the dividing cells at different constriction states. Currently one such mutant exists, Vps4^E209Q^, which causes accumulation of dumbbell-shaped cells in the final stage of division (10). Now, structures of other Cdv proteins allow for rational design of other mutants, such as non-polymerising CdvA or mutations that lock the different ESCRT-IIIs in closed or open states. Observing how these mutants perturb division by live cell imaging, immunofluorescence, or cryo-ET would help to build a more complete model of how these proteins form large ring structures which assemble into, or are remodelled into smaller and smaller radii, leading to abscission. One model to be tested, which is based on previously reported dome-like ESCRT-III assemblies (32, 38), predicts that each additional ring (or rung of spiral) is assembling with decreasing radius relative to the previous rung. Because the division ring constricts over orders of magnitude from ∼1000 nm diameter to around ∼10 nm, several different ESCRT-III proteins may be required (for example B, B1 and B2), able to accommodate the greatly different curvatures experienced by the division ring during this process.

Overall, our data supports a model in which ESCRT-III proteins in archaea operate though the same fundamental principles as their eukaryotic and bacterial counterparts, downstream of archaeal specific CdvA. These ESCRT-III proteins assemble into polymers, engage via their N-terminal amphipathic helices with the membrane, which induces them to undergo conformational transitions important for their function in membrane remodelling.

## Methods

### Bacterial plasmids

**Table 1:** List of protein expression plasmids used.

| Plasmid | Description | Source |
| --- | --- | --- |
| pOPINS SiCdvA | Untagged full-length CdvA from *S. islandicus*. | This work |
| pOPINS SiCdvA<sup>ΔC</sup> | Untagged truncated CdvA from *S. islandicus*, residues 1-205. | This work |
| pOPINS SiCdvA<sup>noF</sup> | Truncated CdvA from *S. islandicus*, residues 1-205, with a C-terminal 7His tag and six mutations that prevent polymerisation (A90D, L94D, I100D, I149D, M153D, I156D). | This work |
| pOPINS SiCdvB | 6His-SUMO-CdvB from *S. islandicus* | This work |
| pOPINS SiCdvB<sup>ΔC I69M I125M</sup> | Truncated 6His-SUMO-CdvB from *S. islandicus*, residues 1-193 and point mutations I69M, I125M. | This work |
| pOPINS SiCdvB1<sup>1-253</sup> | Untagged full-length CdvB1 from *S. islandicus*, residues 1-253. For start codon assignment. | This work |
| pOPINS SiCdvB1<sup>11-253</sup> | Untagged truncated CdvB1 from *S. islandicus*, residues 11-253. For start codon assignment. | This work |
| pOPINS SiCdvB1<sup>46-253</sup> | Untagged truncated CdvB1 from *S. islandicus*, residues 46-253. For start codon assignment. | This work |
| pOPINS SiCdvB1<sup>M46</sup> | 6His-SUMO-CdvB1 from *S. islandicus*, using newly assigned start codon. | This work |
| pOPINS SiCdvB2 | 6His-SUMO-CdvB2 from *S. islandicus*. | This work |
| pOPINSb SaciCdvA | Untagged CdvA of *S. acidocaldarius* | This work |
| pOPINS SaciCdvB | 6His-SUMO-CdvB of *S. acidocaldarius* | This work |
| pOPINS SaciCdvB1<sup>M9</sup> | 6His-SUMO-CdvB1 of *S. acidocaldarius*, residues 9-214 (most likely start codon based on the *S. islandicus* assignment) | This work |
| pOPINS SaciCdvB2 | 6His-SUMO-CdvB2 of *S. acidocaldarius* | This work |
| pOPINS SaciCdvB<sup>Δ15</sup> | 6His-SUMO-CdvB of *S. acidocaldarius*, residues 16-261 | This work |
| pOPINS SaciCdvB1<sup>Δ19</sup> | 6His-SUMO-CdvB1 of *S. acidocaldarius*, with M9 start codon, residues 20-206 (where M9 is residue 1) | This work |
| pOPINS SaciCdvB2<sup>Δ27</sup> | 6His-SUMO-CdvB2 of *S. acidocaldarius*, residues 28-219 | This work |

### Expression plasmid construction

All expression plasmids were constructed in a pOPINS backbone with N-terminal 6His-SUMO. For SiCdvA, SiCdvA^ΔC^, and SaciCdvA constructs, the 6His-SUMO tags were omitted.

Synthetic genes for *S. islandicus* CdvA (AGJ62618.1), CdvB (WP_012711316.1), CdvB1 (WP_014512765.1), CdvB2 (WP_009992298.1), and Vps4 (WP_012711317.1) were ordered from Eurofins. gBlocks for *S. acidocaldarius* CdvA (WP_011278210.1), CdvB (WP_011278209.1), CdvB1 (WP_011277365.1), and CdvB2 (WP_011278248.1) were ordered from Integrated DNA Technologies. All were cloned into a pOPINS backbone using NEBuilder HiFi DNA assembly mix (NEB). Plasmids were transformed into *E. coli* DH5α (Thermo Fisher Scientific) and verified by Sanger sequencing (Eurofins). Point mutations and deletions were introduced by side-directed mutagenesis using NEB’s Q5 Site-Directed Mutagenesis Kit per manufacturer’s protocol.

### SiCdvB1 start codon assignment

Three possible ATG starting positions were identified in the SiCdvB1 sequence (NCBI accession WP_014512765.1). Three expression plasmids of untagged SiCdvB1 were constructed, starting at different positions: M1, M11, and M46. They were transformed into *E. coli* C43(DE3) chemically competent cells. *E. coli* C43(DE3) strains harbouring each of the three SiCdvB1 variants were grown in 100 mL of 2x TY medium with 50 µg/mL kanamycin shaking at 37°C. Protein expression was induced with 1 mM isopropyl-β-D-thiogalactoside (IPTG) and cultured for 4h at 37°C. Cells were harvested by centrifugation.

#### Archaeal growth

*S. islandicus* was cultured in DT medium (39), and S. *acidocaldarius* in Brock medium (40). Both were grown aerobically with stirring, at 75°C and pH 3.0-3.5 until late exponential phase.

#### Western blotting

Crude lysates of *S. islandicus*, *S. acidocaldarius*, and three SiCdvB1 overexpression strains were mixed with SDS loading buffer, heated to 98°C for 5 minutes and ran on a NuPAGE 4-12% Bis-Tris gel (Invitrogen) in MOPS running buffer. Protein was transferred to a nitrocellulose membrane and blocked for an hour with 5% milk and PBS with 0.2% Tween20 (PBST). The membrane was incubated at 4°C overnight with primary α-SaciCdvB1 antibody (10) (1:10,000) in 5% milk and PBST, and then washed three times with PBST for 15 min. This was followed by a 1h incubation in 5% milk and PBST with α-chicken secondary (1:10,000), and a final 15 min PBST wash. The blot was imaged in a Gel Doc XR+ (Bio-Rad, 1.5 s exposure).

### Protein expression and purification

All plasmids were transformed into *E. coli* C43(DE3) cells for protein expression and grown in 2x TY medium supplemented with 50 μg/mL (*S. islandicus* proteins) or 30 μg/mL Kanamycin (*S. acidocaldarius* proteins). After protein expression, cells were collected by centrifugation for 20 min at 4,000*g* at 4°C, flash frozen in liquid nitrogen, and stored at -80°C.

Purification steps were done at room temperature (RT) unless otherwise specified. Cells were lysed either using a cell disruptor at 25 kPsi (Constant Systems) or by sonication (15s on, 10s off, 70% output). Purified proteins were aliquoted, flash-frozen in liquid nitrogen, and stored at -80°C.

Selenomethionine-substituted proteins were also expressed in *E. coli* C43(DE3) as in (41), using a protocol modified from (42). Briefly, feedback inhibition of methionine synthesis was achieved by growing cells in MOPS minimal media and adding amino acids prior to induction with IPTG. For 12 L of culture, the amino acid mix contained 1.2 g of lysine, threonine, phenylalanine, 0.6 g of leucine, isoleucine, valine, and 0.6 g of L-selenomethionine, mixed and portioned to 0.5 g per liter of culture.

#### SiCdvA and SiCdvA^ΔC^

12 L of growth media was inoculated with overnight culture at 1:50 and grown at 15°C, shaking at 200 rpm until optical density OD_600_ ∼0.1. Protein expression was induced with 1 mM IPTG and incubated at 15°C for 4-7 days, reaching OD_600_ ∼6-7.

Cell pellets were dissolved in buffer A1 (30 mM Tris/HCl; 150 mM NaCl; 4 mM TCEP; 0.5% CHAPS (w/v); pH 8.0) with DNAse (Sigma-Aldrich) and 4 cOmplete EDTA-free protease inhibitor tablets (Roche). Cells were lysed by sonication and spun for 15 min at 20,000 *g* at RT to pellet debris. The supernatant and hard pellet were discarded. The wobbly pellet was washed 3-4 times by resuspending in buffer A1 and pelleting at 20,000 *g* for 15 min. CdvA filaments were spun at 15°C for 1 h at 45.000 rpm in a 45Ti rotor (Beckman Coulter) and resuspended in 50 mL of 1 M CHES (pH 9.5) to dissolve filaments. After 30 minutes at RT, the solution was spun at 4°C for 30 min at 35,500 *g* in a JLA16.250 rotor (Beckman Coulter) to remove precipitated protein. Supernatant was dialysed against buffer A1 without CHAPS in 3 kDa Slide-A-Lyzer dialysis cassettes (Thermo Scientific), stirring overnight at RT. The re-formed filaments were spun at 15°C for 3-5 h or overnight at 45,000 rpm and resuspended in the desired volume to reach required concentrations (typically 7.5 mg/mL).

#### SiCdvA^noF^

4 L of growth media was inoculated with 20 mL overnight culture and grown at 15°C, shaking at 200 rpm. Expression was induced with 1 mM IPTG at OD_600_ ∼0.3 and continued at 15°C for 24 h.

Cells were resuspended in 100 mL buffer A2 (30 mM CHES; 100 mM NaCl; 4 mM TCEP; 1% CHAPS (w/v); pH 9.0) and protease inhibitor. The cells were then lysed by sonication and the solution was topped up to 400 mL (final CHAPS concentration 0.25%) and spun at RT for 15 min at 16,250 g to pellet debris. Supernatant was collected and imidazole added to 5 mM, followed by incubation with 10 mL of Super Cobalt NTA affinity resin (Generon) for 45 min. The protein was loaded onto a gravity flow column and washed with buffer A2. Protein was eluted stepwise in buffer A2 with imidazole at concentrations: 25 mM, 50 mM, 75 mM, 100 mM, and 250 mM. Elution fractions containing pure CdvA were pooled, concentrated, and further purified on a HiLoad 16/600 Superdex 200 pg gel filtration column (Cytiva) in buffer A2. Peak fractions were pooled and concentrated to ∼50 mg/mL.

#### SiCdvB, SiCdvB1^M46^, and SiCdvB2

12 L of growth media was inoculated with 60 mL overnight culture and grown at 37°C, shaking at 150 rpm until OD_600_ 0.7-1.0. The temperature was reduced to 18°C and protein expression was induced with 1 mM IPTG, expressing for 20-24 h.

Cells were dissolved in buffer B1 (30 mM Tris/HCl; 150 mM NaCl; pH 8.8), DNAse, and protease inhibitor. Cells were lysed by sonication and centrifuged at 20°C for 20 min at 16,250 *g* to pellet debris. The supernatant was incubated with 20 mL of Super Cobalt NTA resin (Ni-NTA Agarose resin (Qiagen) for SiCdvB) stirring at RT for 25 min. The slurry was loaded on a gravity flow column and washed multiple times with buffer B1 and buffer B1 containing 5 mM imidazole. The protein was eluted 3x with 20 mL buffer B1 with 400 mM imidazole. Elution fractions were pooled in 3 kDa Slide-A-Lyzer dialysis cassettes together with homemade SENP1 (43) and dialysed overnight against buffer B2 (50 mM Tris/HCl; 100 mM NaCl; 4 mM TCEP; pH 8.0). The next day, dialysed sample was incubated in a 70°C water bath for 10 min and immediately spun for 3 min at 16,200 *g* to remove precipitates. This heat-inactivation step was skipped for SiCdvB1^M46^. The supernatant was incubated for 1 h with equilibrated Ni-NTA resin and ran over a gravity column, capturing the flowthrough to collect SiCdvB without affinity tags. 1 M CHES (pH 9.5) was added to a final concentration of 100 mM and the sample was concentrated and further purified on a HiLoad 16/600 Superdex 75 pg gel filtration column (Cytiva) equilibrated with buffer B3 (30 mM CHES; 150 mM NaCl; pH 9.5), and clean fractions concentrated to ∼9.5 mg/mL.

### SiCdvB^ΔC^ ^I69M^ ^I125M^ with selenomethionine

12 L of pre-warmed MOPS media supplemented with 50 μg/mL kanamycin was inoculated with overnight culture and grown at 37°C, 200 rpm for ∼4 h until OD_600_ 0.6-0.7. Then, amino acid mix was added and temperature reduced to 18°C. After 30 min, protein was induced with 1 mM IPTG and expressed overnight before harvesting. The protein was purified as SiCdvB.

#### SaciCdvB, SaciCdvB1, and SaciCdvB2 and their truncations

All *S. acidocaldarius* proteins were expressed by inoculating 12 L of culture 1:100 with overnight culture and growing it to OD_600_ 0.6-0.8, 37°C with shaking at 190 rpm. Protein expression was induced with 0.5 mM IPTG and grown for a further 4 h before harvesting.

Cells were resuspended in buffer B3 with DNase, RNase, and protease inhibitor tablet. They were lysed by sonication and debris pelleted by spinning at 30,000 rpm (Ti45 rotor) for 30 min at 15°C. The supernatant was clarified by filtering through a 0.22 μm filter and imidazole was added to 20 mM. This sample was loaded onto a 5 mL HisTrap HP (Cytiva) equilibrated in buffer B3 with 20 mM imidazole. The column was washed until the UV trace was flat, and then protein was eluted in steps of 50 mM, 100 mM, 200 mM, 300 mM, 500 mM, 1000 mM imidazole. Relevant fractions were pooled and 1 mM TCEP, ∼1 mg homemade GST-tagged SENP1 (43), and 1 mL of pre-washed Glutathione Sepharose 4B resin (Cytiva) were added. All subsequent steps were done at 4°C. The sample was dialysed overnight against buffer B3 with 1 mM TCEP and then ran over a gravity flow column to remove GST-SENP1. To collect cleaved protein, the sample was run over a 5 mL HisTrap HP in buffer B3 with 20 mM imidazole and the flowthrough collected. SaciCdvB was then put through an anion exchange step, which was omitted for SaciCdvB1 and SaciCdvB2. For anion exchange, a 1 mL HiTrap Q HP anion exchange column was equilibrated in buffer B4 (30 mM Tris/HCl; pH 8.9). SaciCdvB was loaded, washed, and gradient eluted to 500 mM NaCl over 35 min. All samples were then subjected to HiLoad 16/600 Superdex 75 pg column in buffer B4 (30 mM CHES; 400 mM NaCl; pH 9.5), and appropriate fractions were pooled and concentrated.

#### Crystal structure determination

Crystallisation conditions were found using our in-house crystallisation facility (44). Many hundreds of commercially available screening conditions were tested for each sample, using vapour-diffusion setups in sitting drop MRC crystallisation plates, mixing 100 nl of sample and 100 nl of reservoir. All crystallisation setups were incubated at 20 °C.

#### SiCdvA^ΔC^

SiCdvA^ΔC^ was best crystallised at 7.5 mg/mL with reservoirs containing 0.282 M KH_2_PO_4_ and 0.094 M Tris/acetate (pH 8.5). For harvesting, crystals were cryoprotected with solutions containing 50 % reservoir solution and 50 % glycerol (v/v). Data was collected on beamline I03 at Diamond Light Source (Harwell, UK). Because the data was weak, partly because of the very long c-axis of the unit cell, data from a number of related crystals, from different conditions, were also collected and subsequently merged into one large dataset with very high multiplicity (26.2, see Table S1 for data collection statistics). Structure determination started with molecular replacement using the PRC subdomain (PDB ID 1PM3). Several copies of the PRC domain could be located using PHASER (45) and the resulting map was improved by NCS averaging using DM as part of the CCP4 package (46). Finally, 14 copies of CdvA could be located and were traced with BUCANEER (47). Manual building in MAIN (48) and refinement with phenix.refine (49) resulted in an atomic model at 2.9 Å resolution (see Table S1 for model and refinement statistics).

#### SiCdvA^ΔCnoF^

SiCdvA^ΔCnoF^ was crystallised with reservoir solution containing 18 % (w/v) PEG 4000, 0.3 M sodium acetate, 0.1 M Tris pH 9.0. Drops to produce diffraction-quality crystals were mixed from 200 nl sample and 200 nl reservoir solution. The sample was at 25 to 50 g/L. Since no additional cryo protection was needed, crystals were harvested directly. Diffraction data was collected on a MarDTB imaging plate detector mounted on an FR-E home rotating anode X-ray source (Rigaku). The structure was solved by molecular replacement (PHASER) using a monomer from the SiCdvA^ΔC^ structure determined above. Manual adjustments in MAIN and refinement with phenix.refine yielded a highly reliable structure at 2.2 Å resolution (see Table S1 for model and refinement statistics).

#### SiCdvB^ΔC I69M I125M^

Because SiCdvB^ΔC^ yielded crystals that diffracted to 3.5 Å, only, which made reliable structure determination difficult, two methionine residues were introduced by mutation, to enable experimental SAD (single wavelength anomalous diffraction) phasing after seleno-methionine labelling: I69 and I125 were mutated to methionine. Using this protein, SiCdvBΔC(I69M, I125M), the best small crystals were obtained using the following reservoir solutions, mixing 100 nl sample at 10 g/L with 100 nl reservoir: 5 % (v/v) 2-propanol, 1M ammonium sulphate or 0.2 M ammonium sulphate, 15 % (w/v) PEG 4000, buffer pH 3.5. The largest crystals were obtained by eventually scaling crystallisation setups to 2 µL sample plus 2 µL reservoir. Crystals were harvested in solutions mixing 30 % (v/v) glycerol with 70 % (v/v) reservoir solution. X-ray diffraction data was collected on beamline I04 at Diamond Light Source (Harwell, UK). Two different crystal forms were solved: P 41 21 2, to 2.7 Å resolution and P 21 21 21, to 2.2 Å resolution. Structure determination used the very good anomalous signal from selenium and phases were obtained using CRANK2 (50). Models were built manually in MAIN and refined with phenix.refine.

### Filament sample preparation and cryo-EM data collection

#### SiCdvA^ΔC^

3 µL of SiCdvA^ΔC^ at 0.05-0.1 mg/mL in buffer A1 was applied to glow-discharged 400-mesh R 2/2 copper grids (Quantifoil), immediately blotted and plunged into liquid ethane using a Vitrobot mark VI (Thermo Fisher Scientific) at RT and 100% humidity. Data was collected on a Titan Krios transmission electron microscope (Thermo Fisher Scientific) operating at 300 keV and equipped with a K2 electron detector (Gatan) in counting mode. EPU software (Thermo Fisher Scientific) was used to automate collection of 2594 micrographs, with a dose of 1 e^-^/Å/s fractionated over 40 frames, nominal pixel size 1.068 Å/px, defocus range of -0.7 to -3.5 μm.

#### SaciCdvB2

3 µL of SaciCdvB2 at 0.4 mg/mL in buffer BB6 (50 mM MES(NaOH); 50 mM NaCl; pH 6.0) was applied to glow-discharged 200-mesh gold R2/2 Quantifoil grids (Quantifoil), blotted and plunged into liquid ethane using a Vitrobot mark VI at 100% humidity and RT. Data was collected on a Titan Krios at 300 keV with a Falcon 4 electron detector (Thermo Fisher Scientific) in electron counting mode. EPU software was used to automate acquisition of 3589 movies, 5 second exposures, total dose 35.86 e^-^/Å^2^ across 40 movie frames at a nominal pixel size of 0.824 Å/px. Defocus ranged between -1.0 to -2.6 µm.

### Helical processing

#### SiCdvA^ΔC^

Movies were motion-corrected using RELION’s implementation of MotionCor2 (51). CTF parameters were estimated with CTFFIND4 (52). A subset of micrographs was manually picked for filaments, and resulting 2D classes used for reference-based autopicking in RELION.

Particles were extracted at 63 Å spacing, with a box of 320 px and 1.068 Å/px. The data underwent rounds of 2D classification, and 253,803 particles were used in 3D refinement. Simulated map from the CdvA^ΔC^ filament solved by X-ray crystallography was used as an initial model. After refining symmetry parameters through 3D refinement, particles underwent CTF refinement and Bayesian polishing. The final reconstruction reached 4.07 Å (twist 48.96°, rise -63.64 Å). The map was sharpened with standard post-processing in RELION.

#### SaciCdvB2

Movies were pre-processed as above. crYOLO was used to train a filament model and pick SaciCdvB2 filaments (53, 54). Helical reconstruction was done in RELION-4.0 (55, 56).

Initial 2D classifications in RELION showed a signal for 19 Å repeats, so boxes were extracted along filaments at 19 Å increments. 3x binned helical segments were extracted with a box size 142 px. Rounds of 2D classification revealed high-twist and low-twist classes. A subset of high-twist and low-twist particles were imported into CryoSPARC (57) for *ab initio* model generation (setting maximum resolution to 8 Å). Helical segments in RELION were then re-extracted at 2x binning (150 px box). Helical symmetry parameters were approximated from *ab initio* models and refined in a 3D classification job. This separated the high-twist (158,229) and low-twist (136,416) particles. Both were first refined via helical reconstruction without symmetry. Particles were again re-extracted at a box size 300 px down-sampled to 200 px and underwent several rounds of helical refinement testing different symmetry parameters. After Bayesian polishing (58) and CTF refinement (59), the low-twist filament structure was refined to 3.2 Å, applying a twist of 1.68°, rise of 38.14 Å, and C2 symmetry. The polished and CTF-refined high-twist particles underwent a round of 3D classification without alignment, which resolved two distinct classes differing in their lumenal densities. High-twist filament class A (139,884 particles) refined to 3.9 Å (twist 176.24°, rise 18.82 Å) and class B (18,345 particles) reached 4.0 Å (twist 176.92°, rise 18.92 Å). All maps were sharpened with standard post-processing in RELION and helical symmetry applied in real-space with relion_helix_tolbox.

### Cryo-EM model building

All cryo-EM volumes and atomic model figures were prepared with ChimeraX (v1.7.1) (60)

#### SiCdvA^ΔC^

A dimer of SiCdvA^ΔC^ from the crystal structure was rigidly docked into the cryo-EM map, and refined in Coot and PHENIX real-space refine.

#### SaciCdvB2

For SaciCdvB2 models, an *ab initio* model was built in the low-twist filament map using ModelAngelo (61). Missing regions were modelled with *Coot* (v.0.9.8.2) (62) and the ISOLDE plugin (63) in ChimeraX (v1.6.1), and real-space refined with PHENIX (49). Several copies of a single chain of this refined model were rigidly docked into the high-twist class A and class B maps. They were re-modelled and refined as above.

### Cryo-EM of lipid nanotubes

#### Lipid nanotube preparation

LNTs were prepared with a modified protocol from (64). Briefly, *E. coli* total lipid extract (Avanti Polar Lipids) dissolved in chloroform and D-galactosyl-β-1,1’ N-nervonoyl-D-erythro-sphingosine (Galactosyl(β) ceramide, Avanti Polar Lipids) in 80:20 chloroform-methanol were mixed together at a 50:50 mass ratio, dried under a gentle stream of nitrogen gas, and left in a desiccator overnight to remove residual solvent. The lipid film was rehydrated in 50 µL of buffer BB6 to a final lipid concentration of 2 mg/mL and incubated at RT for 30 min, then placed on a rotary mixer at RT for 15 min, and finally transferred to a sonication bath for 4 min. Once formed, the LNTs were stored at 4°C.

#### LNT cryo-EM sample preparation and data collection

LNT solution was mixed with various combinations of *Sulfolobus* proteins and incubated for 1 h at 37°C. For most samples, final concentration of LNTs and of each protein component was 0.2 mg/mL, at pH 7. For samples containing Vps4, 2 mM ATP and 4 mM MgCl_2_ were added. For LNT-only grids, their concentration was 1 mg/mL. Grids of each sample were prepared as above, freezing 3µL of solution on an R2/2 200-mesh gold Quantifoil or 400-mesh UltrAuFoil grid in a Vitrobot mark IV. They were imaged in a Glacios at 200 keV with a Falcon III detector.

For the LNT-SaciCdvB2 sample, dataset I was collected on a Titan Krios with a K3, recording 10,599 movies at super-resolution and 2x binning on the fly. A dose of 35.28 e^-^/Å^2^ was delivered over 1.3 s exposures and fractionated over 70 frames, across a defocus range of -1.0 to -2.6 µm. For dataset II, 23,891 movies were collected on a Titan Krios with a K2 detector and BioQuantum energy filter (Gatan). A total dose of 40 e^-^/Å^2^ was fractionated over 43 frames, and a defocus range of -1.2 to -2.7 µm was applied.

#### LNT cryo-EM data processing

Movies were processed with MotionCor2 and CTFFIND4 as above. For dataset I, edges of LNTs were manually picked and used as a basis for template-based autopicking. As there was a clear 38 Å repeat, boxes were extracted with 38 Å spacing. The particles underwent several rounds of 2D classification in Relion. A crYOLO filament model was trained to pick soluble filaments present in this sample. 1.7 million particles were extracted and underwent 2 rounds of 2D classification, 3D refinement without symmetry and 3D classification without alignment. A subset of random 193 thousand particles was refined with applied helical and C2 symmetry, giving a 3.4 Å map of low-twist CdvB2 with comparable helical parameters.

For dataset II, a filament model in crYOLO was trained to pick whole LNTs. Particles were extracted at 38 Å spacing but 2D classification quickly revealed that membrane signal, and not protein lattice signal, was dominating the alignment. To enable alignment of protein signal, the picked coordinates were resampled to 29 Å (out of phase with the 38 Å repeat). A rolling 2D average of these out-of-phase picks enhanced membrane signal – it was then used to subtract membrane signal from the micrographs. The subtraction was implemented as in (27), using scripts for tubulin lattice signal removal (28). 6.5 million crYOLO picks of whole LNTs were extracted from membrane-subtracted micrographs, with a box size of 256 px at 2.73 Å/px. After importing into CryoSPARC particles were split into subsets for faster 2D classification. Initial classification showed presence of two main class types, those with a seam and those without. Data was further classified to separate the two populations and re-imported into Relion for re-extraction at 2.19 Å/px and further 2D classification at finer angular sampling. In the case of classes without a seam, a rare class that showed hints of secondary structure was chosen for further processing. Additional particles were systematically extracted, shifting coordinates along the filament axis at multiples of the 38 Å filament spacing. After more 2D classification and removal of duplicate particles, this increased the total number of quality particles in classes with secondary structure features.

### Pelleting assay

#### S. acidocaldarius lipid extraction

*S. acidocaldarius* was grown as described earlier, harvested and freeze-dried. Archaeal lipid extraction protocol was modified from (65). 4.5 g of cells were extracted in 400 mL of a chloroform-methanol (CHCl_3_-MeOH, 1:1 v/v) mixture in a Soxhlet extractor overnight. The lipid extract was dried and resuspended in 20 mL of H_2_O-MeOH mixture (1:1 v/v) and extensively sonicated for a few hours in a bath sonicator to re-solubilise the extracted lipids. The resulting solution was loaded on a Sep-Pak C18 20 cc Vac cartridge (Waters) used under reduced pressure. The column was washed with 250 mL H_2_O-MeOH (1:1 v/v) and lipids eluted with 250 mL of CHCl_3_-MeOH-H_2_O (1:2.5:1 v/v/v). The solution was dried in a rotary evaporator. The lipids were washed by resuspending in CHCl_3_-MeOH-H_2_O (65:25:4), aliquoting into smaller tubes, and drying in a rotary evaporator. The dry lipids were then weighed and resuspended in CHCl_3_-MeOH-H_2_O.

#### Archaeal liposome preparation

1 mg of *S. acidocaldarius* lipid extract in CHCl_3_-MeOH-H_2_O was dried under a stream of nitrogen gas and left under vacuum overnight to remove residual solvent. The lipid film was then rehydrated with buffer BB6 to 4 mg/mL and shaken vigorously at 80°C for 3h, vortexing every 10-15 minutes. After 8-10 freeze-thaw cycles the liposomes were extruded through a 0.8 µm Whatman membrane filter using a mini-extruder (Avanti) with glass gas-tight syringes (1 mL, Avanti).

#### Pelleting

Proteins were rapidly thawed and resuspended to 0.6 mg/mL in buffer BB6. 20 µL of each sample was centrifuged in a TLA-100 rotor (Beckman Coulter) for 30 min at 100,000 *g* and 4°C to pellet any pre-formed aggregates and polymerised protein. 5 µL of the prespun supernatant was mixed with 30 µL of *S. acidocaldarius* liposomes and incubated at RT for 15 min before spinning for 30 min at 100,000 *g* and 4°C. The pellets were briefly washed and analysed alongside supernatant samples on NuPAGE 4-12% Bis-Tris SDS-PAGE gels (Invitrogen)

### SiCdvB on lipid monolayer

#### Monolayer preparation

Lipid monolayers were prepared on electron microscopy grids using *E. coli* polar lipid extract (Avanti Polar Lipids) (66). A custom Teflon block was filled with 60 μL of buffer (50 mM HEPES(KOH); 100 mM potassium acetate; 5 mM magnesium acetate; pH 7.7). 20 μg of lipids were dissolved in chloroform, applied on top of the buffer, and incubated for 2 min.

Electron microscopy grids (CF300-Cu-UL for negative stain, 300-mesh gold R0.6/1 Quantifoil for cryo-EM) that were baked at 60°C overnight were placed on top of the lipid film, carbon side down, and incubated for 30-60 min. The grids were then carefully lifted, blotted from the side, then washed once with 4 μL of fresh buffer and blotted again.

#### Negative stain sample preparation and imaging

4 μL of SiCdvB at 0.05 mg/mL (diluted with buffer BB6) was applied to monolayer grids, incubated for 30 s, and stained with 2% (w/v) uranyl formate. Grids were imaged on a Tecnai Spirit transmission electron microscope (120 keV, Thermo Fisher Scientific) with a Gatan Orius SC200W camera.

#### Cryo-EM sample preparation, data collection and processing

3 μL of SiCdvB at 0.15 mg/mL (diluted with buffer BB6) was applied to monolayer grids, blotted and plunged into liquid ethane using a Vitrobot mark VI at RT and 100% humidity. Micrographs were collected on a Glacios transmission electron microscope (Thermo Fisher Scientific) operated at 200 keV and equipped with a Falcon III detector in linear mode. Movies, were collected using EPU, using a total dose of 19.7 e^-^/Å^2^ fractionated across 39 movie frames at a nominal pixel size 2.55 Å/px. Defocus ranged between -3 to -5 µm in 0.4 µm increments.

Processing was done in RELION-4.0. Micrographs were motion-corrected with RELION’s implementation of MotionCor2 and CTF estimated with CTFFIND4. Particles were picked with a modified version of Topaz (67, 68). Coordinates were extracted in boxes of 128 pixels and downscaled to 64 pixels for three rounds of 2D classification.

## Acknowledgements

We thank all members of the MRC Laboratory of Molecular Biology (LMB) Electron Microscopy Facility for help and support with data collection and Scientific Computing at the MRC LMB for their computing support. We thank Byung-Gil Lee and Sami Chaaban for their help and discussions relating to cryo-EM data processing. The authors would like to thank Diamond Light Source for beamtime and the staff of beamlines I03 and I04 for assistance with data collection.

This work was funded by the Medical Research Council, UK (U105184326 to J.L.), UK Research and Innovation (MC_UP_1201/27 to B.B.) the Wellcome Trust (227876/Z/23/Z to J.L., 203276/Z/16/Z and 222460/Z/21/Z to B.B.), and the Volkswagen Foundation (94933 to B.B). This study was funded by the Deutsche Forschungsgemeinschaft (DFG, German Research Foundation) under Germany’s Excellence Strategy (CIBSS – EXC-2189 – Project ID 390939984).

For the purpose of open access, the MRC Laboratory of Molecular Biology has applied a CC BY public copyright licence to any author accepted manuscript version arising.

