## Supporting figures and tables for "Molecular structure of the ESCRT III-based archaeal CdvAB cell division machinery"


**This file includes:**

Figures S1 to S4

Tables S1 to S3

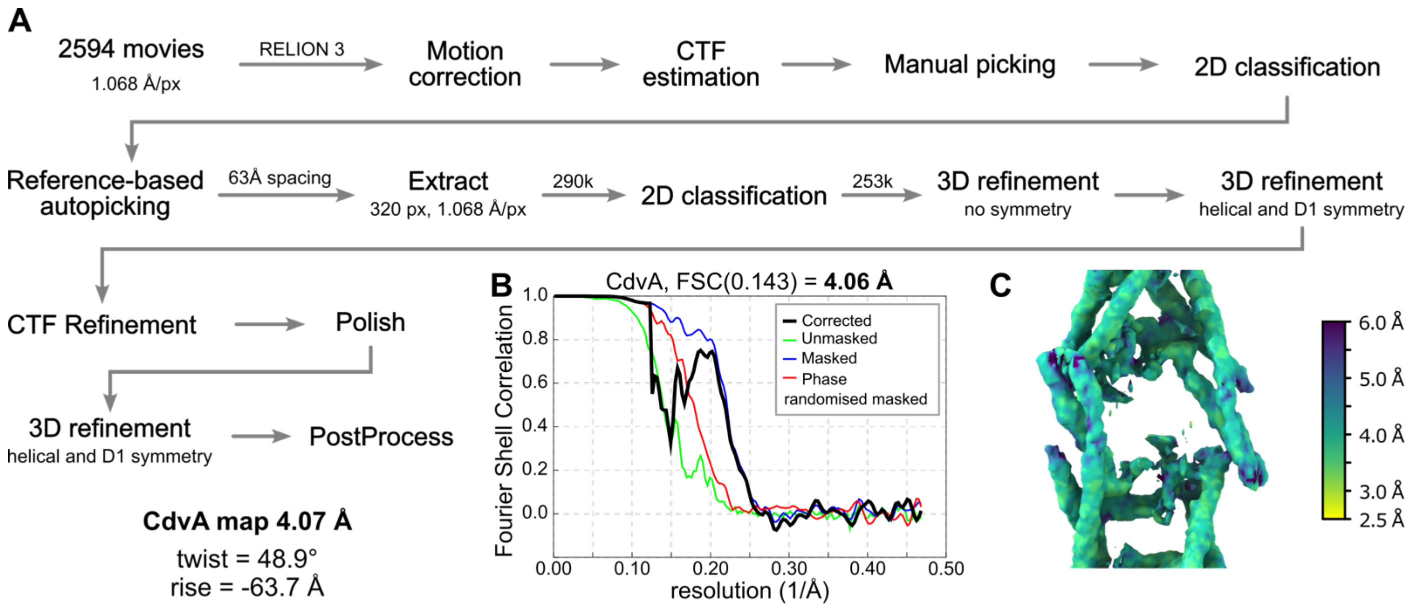


**Figure S1: CryoEM of *S. islandicus* CdvA$^{\Delta C}$.**

**(A)** Cryo-EM processing flowchart for the *S. islandicus* CdvA$^{\Delta C}$ filament structure. All steps were performed in Relion. **(B)** FSC plot and **(C)** local resolution estimations of the CdvA$^{\Delta C}$ filament.

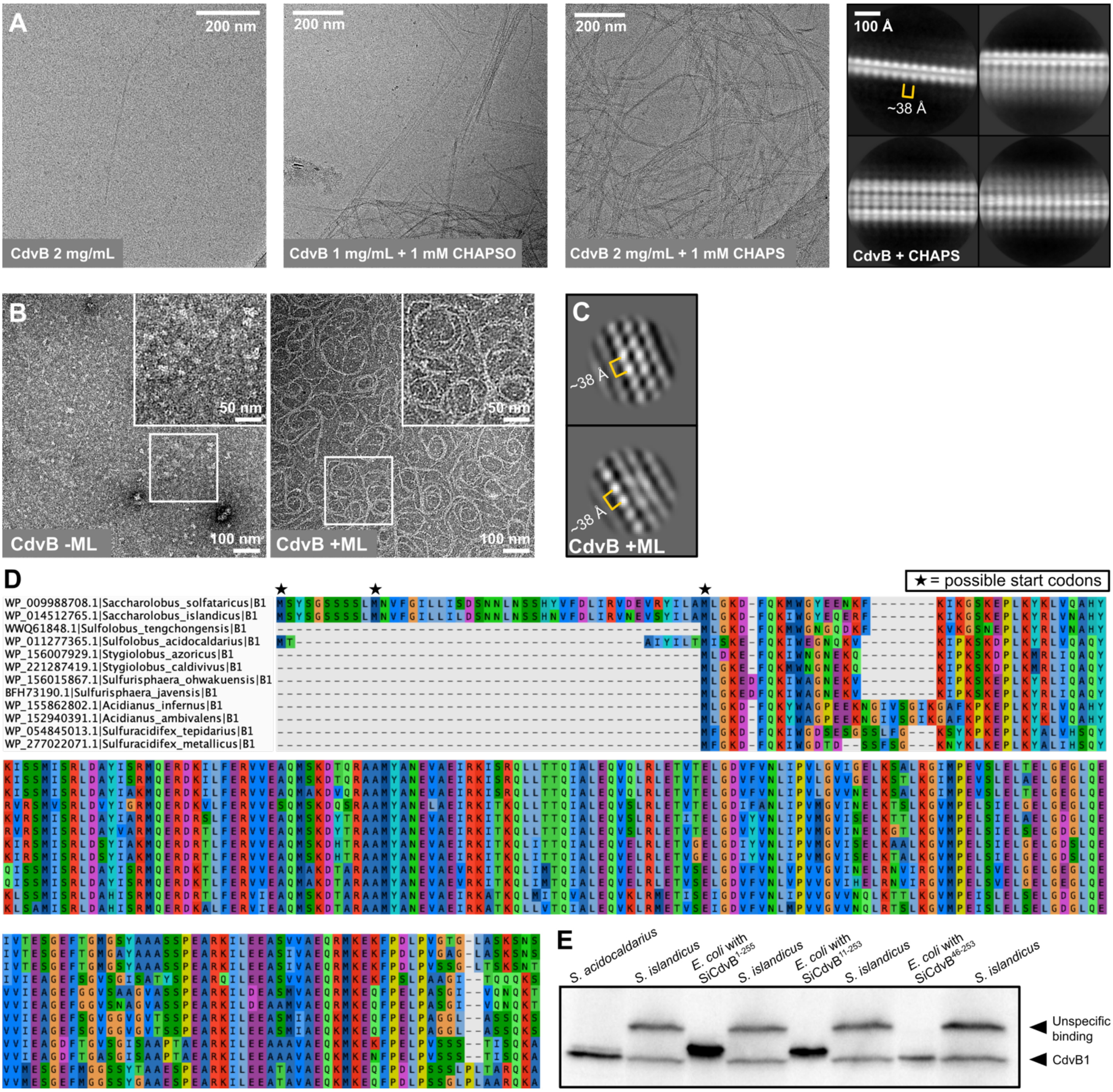


**Figure S2: Polymerisation of CdvB and start codon assignment of CdvB1.**

**(A)** Cryo-EM images of *S. islandicus* CdvB filaments and bundles under different buffer conditions. **(B)** Negative stain electron micrographs of CdvB in absence (left) and presence (right) of a lipid monolayer of *E. coli* polar lipid extract. **(C)** CryoEM 2D class averages of CdvB on lipid monolayer show ~38 Å subunit spacing **(D)** Sequence alignment of full-length *Sulfolobales* CdvB1 sequences from the NCBI protein database. **(E)** Western Blot of CdvB1 (α-SaciCdvB1 antibody) on *S. aciodocaldarius* and *S. islandicus* cell lysates, alongside *E. coli*-expressed SiCdvB1 constructs with different start codon positions. The *S. acidocaldarius* CdvB1 antibody recognises *S. islandicus* CdvB1 but also produces an unspecific signal at a higher molecular weight.

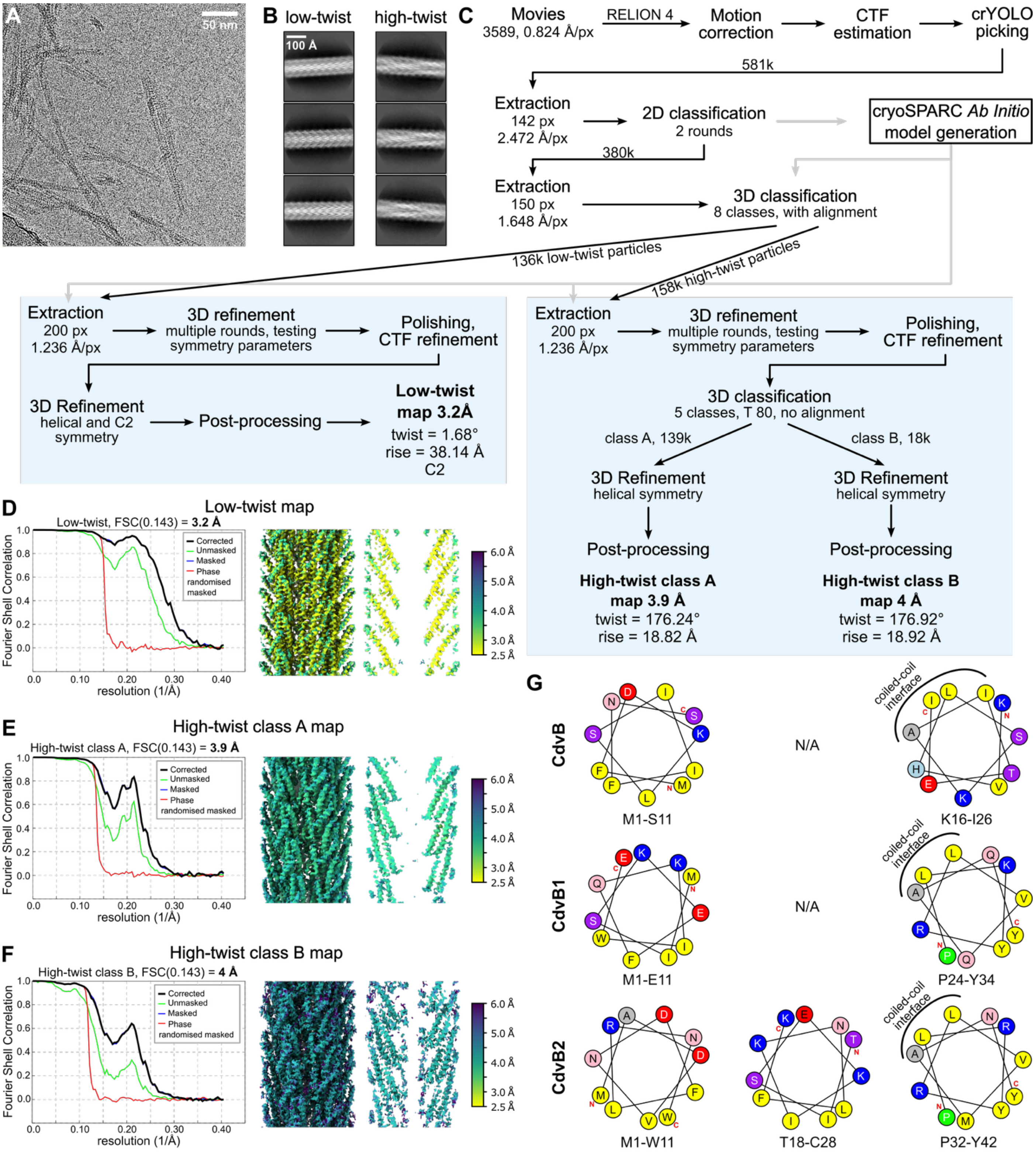


**Figure S3: Cryo-EM processing of CdvB2 filaments.**

**(A)** Representative cryo-EM micrograph of *S. acidocaldarius* CdvB2 filaments. **(B)** Example 2D class averages of high-twist and low-twist CdvB2 filaments. **(C)** Cryo-EM processing flowchart for obtaining three CdvB2 cryo-EM maps. All steps except cryoSPARC *ab initio* model generation were performed in Relion 4. **(D, E, F)** FSC plots (left) and local resolution maps (right) of the three CdvB2 maps. **(G)** Helical wheel diagrams of *S. acidocaldarius* CdvB and CdvB1 N-terminal regions that correspond to amphipathic helices of CdvB2 (shown in Figure 3H-K).

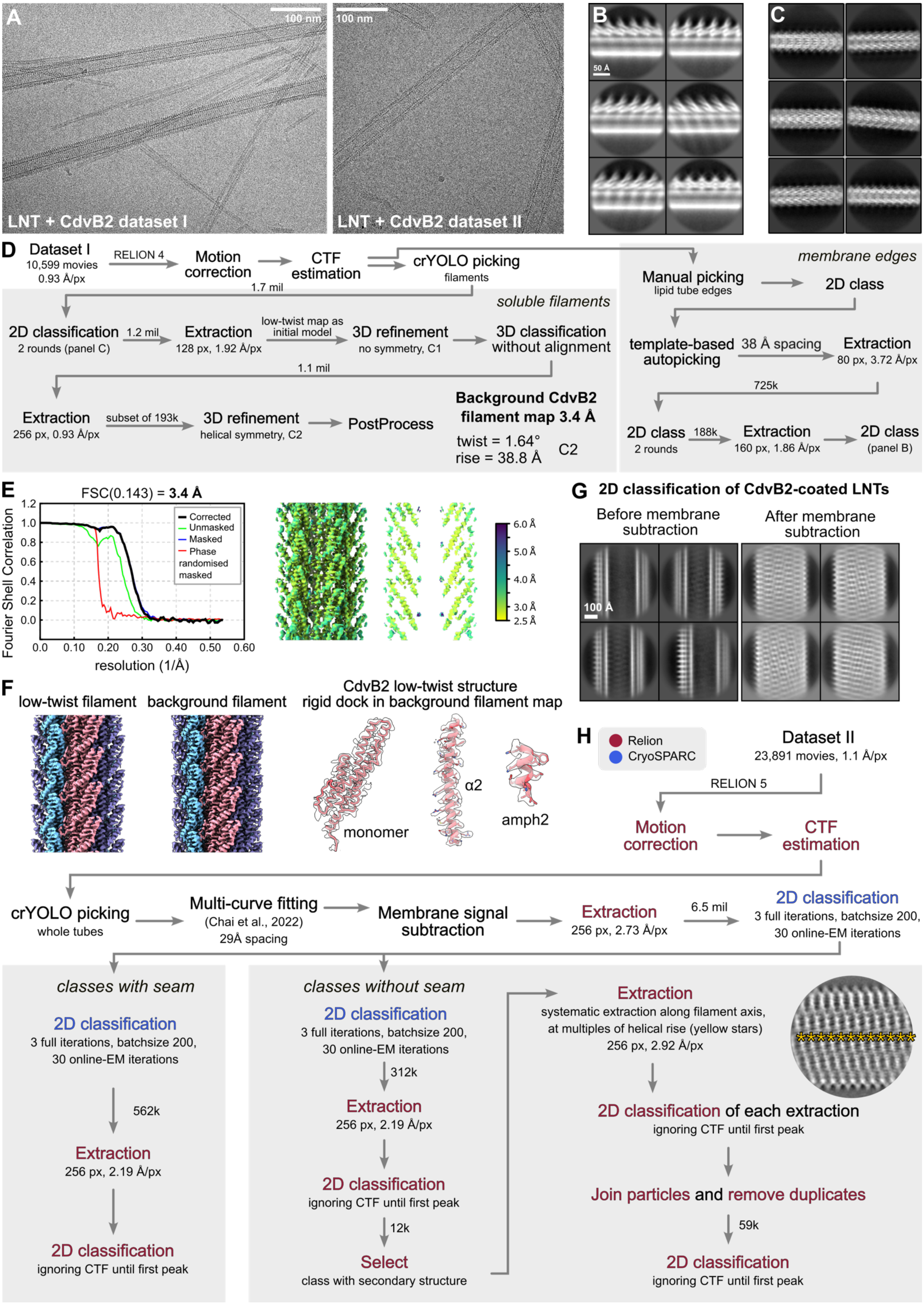

A
100 nm
LNT + CdvB2 dataset I
100 nm
LNT + CdvB2 dataset II
B
50 Å
C
D
Dataset I
10,599 movies
0.93 Å/px
RELION 4
Motion correction
CTF estimation
1.7 mil
crYOLO picking
filaments
soluble filaments
2D classification
2 rounds (panel C)
1.2 mil
Extraction
128 px, 1.92 Å/px
low-twist map as initial model
3D refinement
no symmetry, C1
3D classification without alignment
1.1 mil
Extraction
256 px, 0.93 Å/px
subset of 193k
3D refinement
helical symmetry, C2
PostProcess
Background CdvB2 filament map 3.4 Å
twist = 1.64°
rise = 38.8 Å
C2
membrane edges
Manual picking
lipid tube edges
2D class
template-based autopicking
38 Å spacing
Extraction
80 px, 3.72 Å/px
725k
2D class
2 rounds
188k
Extraction
160 px, 1.86 Å/px
2D class
(panel B)
E
FSC(0.143) = 3.4 Å
Fourier Shell Correlation
resolution (1/Å)
Corrected
Unmasked
Masked
Phase randomised masked
6.0 Å
5.0 Å
4.0 Å
3.0 Å
2.5 Å
F
low-twist filament
background filament
CdvB2 low-twist structure rigid dock in background filament map
monomer
α2
amph2
G
2D classification of CdvB2-coated LNTs
Before membrane subtraction
After membrane subtraction
100 Å
H
Relion
CryoSPARC
Dataset II
23,891 movies, 1.1 Å/px
RELION 5
Motion correction
CTF estimation
crYOLO picking
whole tubes
Multi-curve fitting
(Chai et al., 2022)
29Å spacing
Membrane signal subtraction
Extraction
256 px, 2.73 Å/px
6.5 mil
2D classification
3 full iterations, batchsize 200, 30 online-EM iterations
classes with seam
2D classification
3 full iterations, batchsize 200, 30 online-EM iterations
562k
Extraction
256 px, 2.19 Å/px
2D classification
ignoring CTF until first peak
classes without seam
2D classification
3 full iterations, batchsize 200, 30 online-EM iterations
312k
Extraction
256 px, 2.19 Å/px
2D classification
ignoring CTF until first peak
12k
Select
class with secondary structure
Extraction
systematic extraction along filament axis, at multiples of helical rise (yellow stars)
256 px, 2.92 Å/px
2D classification of each extraction
ignoring CTF until first peak
Join particles and remove duplicates
59k
2D classification
ignoring CTF until first peak

**Figure S4: Cryo-EM of lipid nanotubes (LNTs) coated with *S. acidocaldarius* CdvB2.**

**(A)** Example micrographs of two collected datasets of LNTs and CdvB2. **(B)** 2D class averages of the edges of LNT membranes coated with CdvB2 in dataset I. **(C)** 2D class averages of CdvB2-only filaments in dataset I. **(D)** Cryo-EM processing pipeline for soluble background CdvB2 filaments and CdvB2-coated LNTs in dataset I. All steps were performed in Relion 5 unless specified. **(E)** FSC plot (left) and local resolution map (right) of the soluble background filament map. **(F)** Low-twist CdvB2 filament map from Figure 3 (left) and the soluble background filament map (middle). The modelled low-twist structure rigidly docked into the background filament map (left). **(G)** Example 2D class averages before (left) and after (right) subtraction of the dominating membrane signal. **(H)** Cryo-EM processing pipeline for CdvB2-coated LNTs in dataset II. Relion jobs are shown in red, CryoSPARC jobs in blue.

### Table S1: X-ray Crystallography data collection and refinement statistics

| | ***S. islandicus*** **CdvA$^{\Delta C}$** (PDB 9S9G) | ***S. islandicus*** **CdvA$^{\Delta CnoF}$** (PDB 9S9I) | ***S. islandicus*** **CdvB$^{\Delta C}$ (I69M I125M) closed** (PDB 9S9J) | ***S. islandicus*** **CdvB$^{\Delta C}$ (I69M I125M) semi open** (PDB 9S9K) |
|---|---|---|---|---|
| **Data collection and processing** | | | | |
| Space group | $P2_1$ | C2 | $P\ 4_1\ 2_1\ 2$ | $P\ 2_1\ 2_1\ 2_1$ |
| Wavelength (Å) | 0.97953 | 1.54179 | 0.97942 | 0.97942 |
| Beamline | I03, Diamond Light Source | In-house | I04, Diamond Light Source | I04, Diamond Light Source |
| *a*, *b*, *c* (Å)<br>α, β, γ (°) | 71.03, 102.72, 328.55, 90.00, 92.35, 90.00 | 66.33, 78.68, 93.28, 90.00, 104.63, 90.00 | 59.35, 59.35, 124.37, 90.00, 90.00, 90.00 | 54.72, 73.34, 97.68, 90.00, 90.00, 90.00 |
| Resolution (Å) | 2.9 | 2.2 | 2.7 | 2.2 |
| $R_{meas}$ / $R_{pim}$ | 1.711 / 0.358 | 0.077 (0.864) | 0.051 (1.129) | 0.093 (0.890) |
| CC1/2 | 0.672 (0.137) | 0.998 (0.727) | 1.0 (0.829) | 0.999 (0.779) |
| I/σ(I) | 1.5 (0.2) | 9.7 (1.5) | 23.8 (2.4) | 11.0 (1.9) |
| Completeness (%) | 98.5 | 96.5 (91.5) | 100 (100) | 99.9 (99.9) |
| Multiplicity | 26.2 | 3.0 | 12.4 | 6.5 (6.8) |
| **Refinement** | | | | |
| Phasing | Molecular replacement: PDB 1PM3 | Molecular replacement: PDB 9S9G (this work) | SeMet SAD phasing | SeMet SAD phasing |
| Molecules per ASU | 14 | 2 | 1 | 2 |
| Number of reflections | 36,663 | 22,832 | 6,575 | 20,539 |
| $R_{work}$ / $R_{free}$ | 0.226 / 0.298 | 0.228 / 0.259 | 0.243 / 0.266 | 0.223 / 0.277 |
| *Model composition* | | | | |
| Non-hydrogen atoms | 21,978 | 3,188 | 1,352 | 2,856 |
| Protein residues | 2,711 | 384 | 171 | 352 |
| Ligands | 0 | 0 | 0 | 0 |
| *B factors (Å$^2$)* | | | | |
| Protein | 68.90 | 63.43 | 101.15 | 57.15 |
| *R.m.s. deviations* | | | | |
| Bond lengths (Å) | 0.011 | 0.010 | 0.007 | 0.004 |
| Bond angles (°) | 1.432 | 1.353 | 0.950 | 0.893 |
| *Validation* | | | | |
| MolProbity score | 2.2 | 1.64 | 1.99 | 1.78 |
| Clashscore | 16.66 | 13.43 | 15.68 | 7.68 |
| Poor rotamers (%) | 4.29 | 0.28 | 0.00 | 3.00 |
| *Ramachandran plot* | | | | |
| Favoured (%) | 98.4 | 98.15 | 95.81 | 98.55 |
| Disallowed (%) | 0.04 | 0.00 | 0.00 | 0.00 |

Statistics for the highest-resolution shell are shown in parentheses.

### Table S2: Cryo-EM data collection, refinement and validation statistics

| | ***S. islandicus* CdvA$^{\Delta C}$** (EMD-54678) (PDB 9S9H) | ***S. acidocaldarius* CdvB2 low-twist** (EMD-54673) (PDB 9S97) | ***S. acidocaldarius* CdvB2 high-twist class A** (EMDB-54674) (PDB 9S98) | ***S. acidocaldarius* CdvB2 high-twist class B** (EMDB-54675) (PDB 9S99) |
|---|---|---|---|---|
| **Data collection and processing** | | | | |
| Magnification | 75,000 | 96,000 | 96,000 | 96,000 |
| Voltage (kV) | 300 | 300 | 300 | 300 |
| Electron exposure (e–/Å$^2$) | 40 | 35.86 | 35.86 | 35.86 |
| Defocus range (μm) | -0.7 to -3.5 | -1.0 to -2.6 | -1.0 to -2.6 | -1.0 to -2.6 |
| Pixel size (Å) | 1.068 | 0.824 | 0.824 | 0.824 |
| Symmetry imposed | D1, 48.92° twist, -63.70 Å rise | C2, 1.68° twist, 38.14 Å rise | 176.24° twist, 18.82 Å rise | 176.92° twist, 18.92 Å rise |
| Initial particle images (no.) | 290,032 | 580,965 | 580,965 | 580,965 |
| Final particle images (no.) | 253,803 | 136,416 | 139,884 | 18,345 |
| Map resolution (Å) | 4.1 | 3.2 | 3.9 | 4.0 |
| FSC threshold | 0.143 | 0.143 | 0.143 | 0.143 |
| **Refinement** | | | | |
| Initial model used (PDB code) | 9S9G (this work) | *De novo* (with ModelAngelo) | *De novo* (with ModelAngelo) | *De novo* (with ModelAngelo) |
| Map sharpening *B* factor (Å$^2$) | -139 | -83 | -125 | -90 |
| *Model composition* | | | | |
| Non-hydrogen atoms | 3130 | 4549 | 4530 | 4728 |
| Protein residues | 386 | 571 | 567 | 592 |
| Ligands | 0 | 0 | 0 | 0 |
| *B factors (Å$^2$)* | | | | |
| Protein | | | | |
| *R.m.s. deviations* | | | | |
| Bond lengths (Å) | 0.005 | 0.004 | 0.005 | 0.005 |
| Bond angles (°) | 0.618 | 0.896 | 1.05 | 0.988 |
| *Validation* | | | | |
| MolProbity score | 1.93 | 1.28 | 1.27 | 1.42 |
| Clashscore | 15.25 | 5.17 | 5.08 | 6.65 |
| Poor rotamers (%) | 0.57 | 0.20 | 0.00 | 0.19 |
| *Ramachandran plot* | | | | |
| Favored (%) | 96.34 | 99.82 | 99.11 | 97.77 |
| Allowed (%) | 3.66 | 0.18 | 0.89 | 2.23 |
| Disallowed (%) | 0.00 | 0.00 | 0.00 | 0.00 |

### Table S3: Structures summary table

<table>
<tr><td></td><td colspan="2">S. islandicus CdvA$^{\Delta C}$</b></td><td><b><i>S. islandicus</i> CdvA$^{\Delta CnoF}$</b></td><td colspan="2"><b><i>S. islandicus</i> CdvB$^{\Delta C\ I69M\ I125M}$</td><td colspan="3">S. acidocaldarius CdvB2</td></tr>
<tr><td>NCBI Protein ID</td><td colspan="2">AGJ62618.1</td><td>AGJ62618.1</td><td colspan="2">WP_012711316.1</td><td colspan="3">WP_011278248.1</td></tr>
<tr><td>Sequence used in experiment<br><br>mutations are underlined in bold, truncated regions are in grey italics</td><td colspan="2">MPVSYEVLTKFIGQKVKDIYGREF<br>GYLIHVYSEIDGSITGIEVAQGSSIL<br>TMGPERIKLDGDSILILPDWKAEAI<br>RILSLMEKIRKRQRALEELYNKQEI<br>PKSDYDDMKRKLDTEMLKVKDDQ<br>NKLKGKLKSRLNDIEDQLAHIDKAV<br>ISLKMSYISSEIPENAYKGSMEVLR<br>QSKDSYTLERDDIRKTLDRLDSLD<br>KESIELKPLGSLSTSQQGEAKSDQ<br>SKSEIPLPIPVKVINTL</td><td>MPVSYEVLTKFIGQKVKDIYGREFG<br>YLIHVYSEIDGSITGIEVAQGSSILTM<br>GPERIKLDGDSILILPDWKAEAIRILS<br>LMEKIRKRQR<u>D</u>LEE<u>D</u>YNKQE<u>D</u>PKSD<br>YDDMKRKLDTEMLKVKDDQNKLKG<br>KLKSRLNDIEDQLAHIDKAV<u>D</u>SLK<u>D</u>S<br>Y<u>D</u>SSEIPENAYKGSMEVLRQSKDSY<br>TLERDDIRKTLDRLDSLDKESIELKPL<br>GSLSTSQQGEAKSDQSKSEIPLPIPV<br>KVINTL</td><td colspan="2">MFDKLPFIFNNEKRRKAQLGKILTEISLK<br>LKDQQTRLEEAIRRLKDRDKELFEKVVR<br>AQVEGDDAKAK<u>M</u>YAQEIADIRRIIKVIYT<br>AFLAIEKVRLKLDTVQELQGVSLVLYPV<br>AKILGDLKDQ<u>M</u>KGIAPEVAIALDSIISSVN<br>GIAVETGAINDRGVVPAVVDEQARQILD<br>EAQKMAEVKVRELLPDLPHPPIEQSSR<br>VSQSRPAVRKITERELLDYIVNNGGFLDI<br>EHFSKVYGVEKQEVVKLLEVLKSKGLIA<br>VES</td><td colspan="3">MADVNDFLRNWGGRQEPTISEKIKNLFKSQ<br>QPLRYRLVMANYRLRTTISRLDVYISKLQERD<br>RSLFEKVVESQISKDSARAAMYANEIAEIRKIT<br>KQLLTTEIALEQVQLRLETITEIGDIFTSLVPVI<br>GVIRELRNVMKGVMPELSIELADLEEGLQEV<br>VLEAGEFTGARVDFATSSPEARKILDEASAV<br>AEQRMKEKFPSLPSFATSVDQKTNANQK</td></tr>
<tr><td>Method</td><td>X-ray crystallography</td><td>Cryo-EM</td><td>X-ray crystallography</td><td colspan="2">X-ray crystallography</td><td colspan="3">Cryo-EM</td></tr>
<tr><td>Resolution (Å)</td><td>2.9</td><td>4.1</td><td>2.2</td><td>Closed form: 2.7</td><td>Semi-open form: 2.2</td><td>Low-twist: 3.2</td><td>High-twist class A: 3.9</td><td>High-twist class B: 4.0</td></tr>
<tr><td>PDB ID</td><td>9S9G</td><td>9S9H</td><td>9S9I</td><td>9S9J</td><td>9S9K</td><td>9S97</td><td>9S98</td><td>9S99</td></tr>
</table>
